# Organelle bridges and nanodomain partitioning govern targeting of membrane-embedded proteins to lipid droplets

**DOI:** 10.1101/2024.08.27.610018

**Authors:** Arda Mizrak, Jacob Kæstel-Hansen, Jessica Matthias, J. Wade Harper, Nikos S. Hatzakis, Robert V. Farese, Tobias C. Walther

## Abstract

Numerous metabolic enzymes translocate from the ER membrane bilayer to the lipid droplet (LD) monolayer, where they perform essential functions. Mislocalization of certain LD-targeted membrane proteins, including HSD17B13 and PNPLA3, is implicated in metabolic dysfunction-associated steatotic liver disease (MASLD). However, the mechanisms governing the trafficking and accumulation of ER proteins on LDs remain poorly understood. Here, using MINFLUX and HILO single-molecule tracking combined with machine learning, we show that HSD17B13, GPAT4, and the model cargo *LiveDrop* diffuse at comparable speeds in the ER and on LDs, but become nano-confined upon reaching the LD surface. Mechanistic dissection of *LiveDrop* targeting revealed that this confinement, along with protein accumulation on LDs, depends on specific residues within its targeting motif. These residues mediate preferential and repeated interactions with nanoscale membrane domains, suggesting that LD-targeted proteins selectively partition into distinct lipid-protein environments that transiently retain and concentrate them at the LD surface. Single-molecule trajectories further revealed bidirectional trafficking of *LiveDrop* across seipin-containing ER-LD bridges, providing direct evidence for lateral protein transfer across membrane contact sites. These findings establish nanodomain-based confinement as a key mechanism driving selective protein accumulation on LDs and reveal how membrane bridges between organelles facilitate protein sorting.

---

Lipid droplets (LDs) serve as hubs for cellular metabolism. Unlike membrane-bound organelles, LDs are composed of a hydrophobic core of neutral lipids, primarily triacylglycerols (TGs) and cholesterol esters (CEs), surrounded by a phospholipid monolayer, rather than a bilayer membrane. This unique architecture supports a specialized proteome rich in enzymes that coordinate lipid synthesis and mobilization. In addition to their established roles in energy storage and membrane production, LD-associated proteins play critical roles in maintaining metabolic homeostasis. Notably, proteins, such as PNPLA3 and HSD17B13, have been implicated in the pathogenesis of metabolic disorders, including metabolic dysfunction-associated steatotic liver disease (MASLD)^1–5^.

Numerous LD proteins target to the LD surface from the ER bilayer membrane. These proteins insert into the ER membrane during their synthesis and subsequently traffic to LDs, where they favor localization and accumulation at the monolayer surface. How LD proteins transition between the two organelles in human cells is largely unknown. A current model posits that they traffic to LDs through ER-LD membrane bridges^6–9^. It remains unclear whether individual LDs are connected to the ER through multiple membrane bridges, as observed in *Drosophila* cells ^7^, whether the bilayer–monolayer attachment at ER–LD contact sites is structurally stable, or how this unique membrane topology influences protein mobility between the two compartments. Also unclear is how ER-derived cargoes accumulate at LDs, rather than equilibrating between the LD and ER. Possible explanations to drive cargo accumulation on LDs include a change in conformation, an LD trapping mechanism, or selective ER protein degradation^8,10,11^.

Analyzing ER-LD cargo trafficking is challenging because the transition between ER and LDs is extremely rapid, and hence, these events are rare in the lifetime of a protein. Additionally, ER to LD trafficking occurs within a dense ER network that maintains contact sites with LDs and other organelles, making events hard to track. Here, we sought to overcome these challenges in capturing protein targeting between ER and LD by employing single-molecule tracking with MINFLUX and HILO fluorescence microscopy.

## ER-to-LD cargoes move with similar speeds in the ER and at LDs, but are nano-confined at LDs

To analyze ER-LD membrane protein trafficking, we expressed and sparsely labeled different LD protein cargoes (GPAT4 and HSD17B13) fused to a Halo tag with fluorescent dyes in human SUM159 cells, which display high lipid storage capacity^12,13^. This allowed for tracking single molecules in live cells using minimal fluorescence photon fluxes (MINFLUX) nanoscopy^14,15^.

We recorded a total of 2 million single-molecule localizations with 3.5-kHz sampling rate and an estimated <15-nm spatial resolution in the x-y axes (Extended Fig. 1a, b). To analyze the datasets for each cargo, we created a pipeline that first classifies each single-molecule track based on its subcellular location using BODIPY-positive LDs as fiducial markers (Extended Data Fig. 1c, d). We then calculated the apparent diffusion coefficients for each compartment using a 5-ms rolling window mean-squared displacement (MSD). GPAT4’s speed of motion, reflected in the value of the jump distance and its apparent diffusion constant D_app_, was unaffected by the transition of the molecule from the ER bilayer to the LD monolayer, with similar D_app_ for both locations (LD_(median)_ 0.078 µm^2^/s, ER_(median)_ 0.085 µm^2^/s) (Fig. 1c, Extended Data Fig. 1e). Similarly, HSD17B13 showed comparable speeds on both membranes (Fig. 1c, Extended Data Fig. 1e). The similar molecular speeds of each protein in both the ER and LD compartments suggests similar fluidity and membrane crowding in the compartments.

**Fig. 1.**
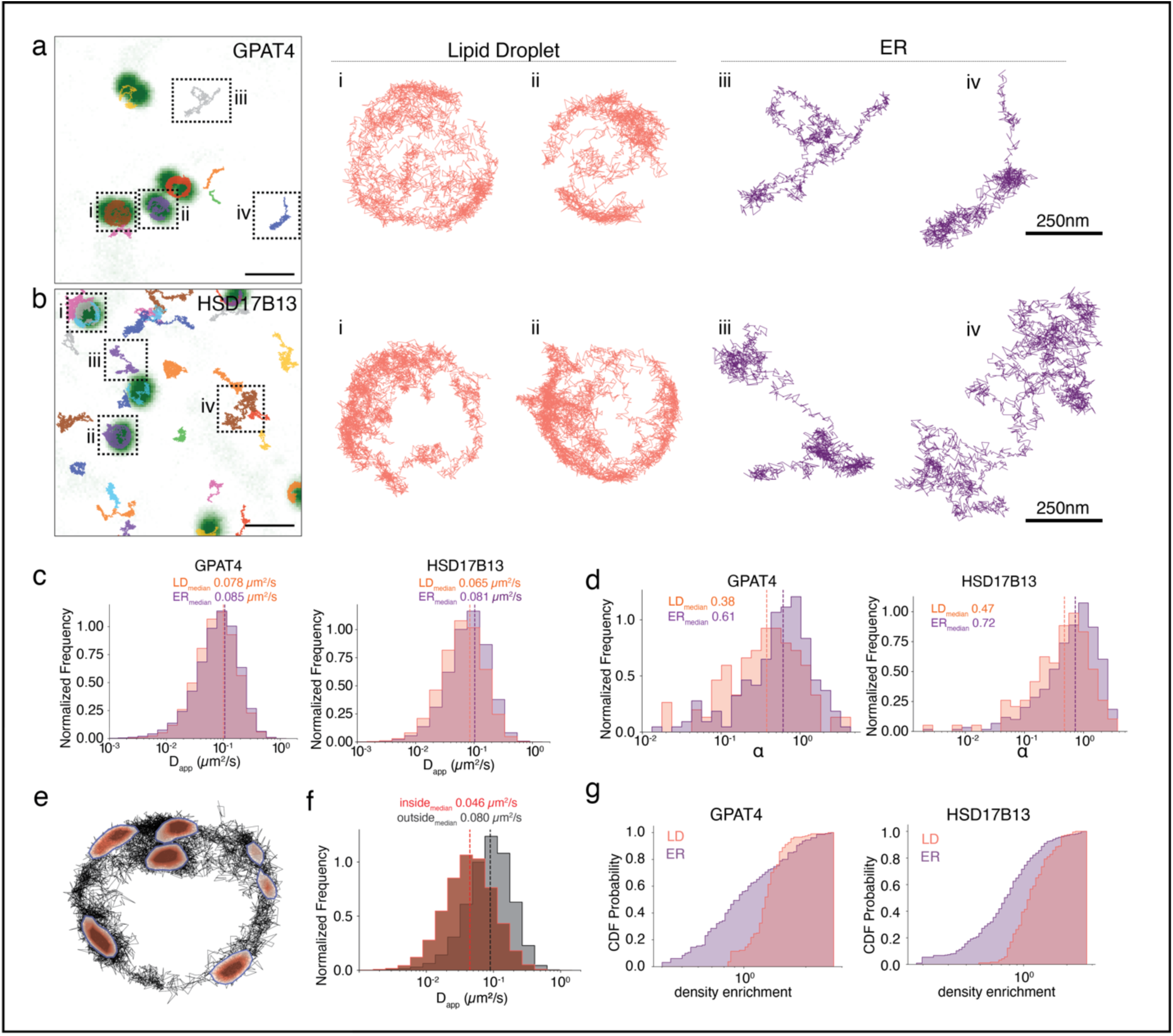
Nanodomain interactions alter molecular motion on LD surface. **a, b**. MINFLUX single-molecule trajectories of GPAT4 (a) and HSD17B13 (b) overlaid on confocal images of cells stained with BODIPY 493/503 to label LDs. Confocal LD images were used to classify trajectories by subcellular localization. Representative examples of single-molecule tracks on LDs and the ER are shown on the right. Scale bars: 1 µm (left), 250 nm (right). **c.** Diffusion coefficients for GPAT4 (left) and HSD17B13 (right), calculated using a 5-ms rolling-window mean squared displacement (MSD) analysis. Values are shown separately for ER-localized (purple) and LD-localized (salmon) trajectories. Dotted lines indicate medians. *N*_GPAT4_ = 301, *N*_HSD17B13_ = 785. **d.** Histograms of the anomalous diffusion exponent (α) for ER-localized (purple) and LD-localized (salmon) tracks of GPAT4 (left) and HSD17B13 (right). α values were obtained by fitting a power-law equation to the full MSD curve of each trajectory. Dotted lines indicate medians. **e.** Kernel density estimate (KDE) analysis of GPAT4 single-molecule trajectories on LDs. Tracks were segmented into 100 ms intervals, and localization density was computed using a Gaussian kernel. Dark red contours indicate regions of elevated localization density, representing putative nanodomains on the LD phospholipid monolayer. **f.** Distribution of the apparent diffusion coefficient (D_app_) of LD cargo within (red) or outside (gray) nanodomain regions on LDs. D_app_ was calculated using a 5-ms rolling window mean square displacement (MSD) analysis and classified based on the molecule’s position relative to nanodomain boundaries. N = 2155. **g.** ER and LD membrane for GPAT4 and HSD17B13 were plotted on a cumulative distribution function graph. Density enrichment was calculated by driving the ratio of mean KDE density at each dense spot and total grid normalized KDE density.

We next tested for anomalous diffusion by computing MSD curves across the full duration of single-molecule tracks on the ER or LD membrane. These full-length MSD curves were then fit to the power-law model MSD = 4 D_app_ t^α^, where α≠ 1 indicates deviation from ideal Brownian motion. Unexpectedly, despite similar D_app_, we observed differences in the temporal correlations of molecular motion between the ER and LDs. On LDs, cargos exhibited lower α, reflecting sub-diffusive behavior, where the MSD was not linearly proportional to the time passed, and instead, there was a power law effect (Extended Fig. 1f). Correspondingly, the median α -value for GPAT4 and HSD17B13 tracks were lower on LDs (0.38 and 0.47, respectively) than at the ER (0.61 and 0.72, respectively) (Fig. 1d).

The more anomalous diffusion of GPAT4 and HSD17B13 at LDs suggest that these cargoes may be dynamically confined to nano-domains on LDs despite their similar movement speed. To address this possibility, we mapped the nano-scale distribution of cargoes in the ER and LDs. We segmented individual single-molecule trajectories into 100-ms windows and quantified spatial localization density using Gaussian kernel density estimation (KDE) across ER and LD membranes (Fig. 1e, Extended Data Fig. 2a). We observed small regions (0.027 ± 0.013 µm^2^) of dense cargo localizations in both compartments where D_app_ decreased by 40% upon entry (Fig 1f, Extended Data Fig. 2b). Although these nanodomains were observed in both the ER and LD membranes, cargo localized to LDs exhibited significantly higher localization densities within these regions, despite lower normalized total density across the entire track (Extended Data Fig. 2c–e). This resulted in a higher density enrichment on LDs, indicating that proteins preferentially accumulate within spatially restricted domains rather than being evenly distributed (Fig. 1g). Based on time-resolved KDE segmentation, we found that cargo on LDs exhibited significantly more frequent transitions into and out of nanodomains than the ER localized proteins, suggesting dynamic and repeated sampling of these LD membrane subregions (Extended Data Fig. 2f). These findings suggest that cargo exhibits frequent and spatially concentrated interactions with nanodomains on LD monolayer, which may contribute to the lower anomalous exponent (α) observed at LDs.

## The change in protein behavior on LDs is mediated by specific amino acid residues

LD proteins, such as GPAT4 or HSD17B13, often contain multiple regions that can bind LDs ^1,6,16^, complicating the analysis of the underlying mechanisms for membrane protein targeting. To reduce the complexity of the system, we utilized a central hydrophobic hairpin region of GPAT4, known as *LiveDrop,* which is sufficient to recapitulate ER-to-LD targeting and LD accumulation^6,8^. *LiveDrop* recapitulated molecular motions of GPAT4 and HSD17B13, showing similar molecular speeds in both compartments and anomalous diffusion with nano-confinement at LD surfaces (Fig. 2a-c Extended Fig. 3b-f).

**Fig. 2.**
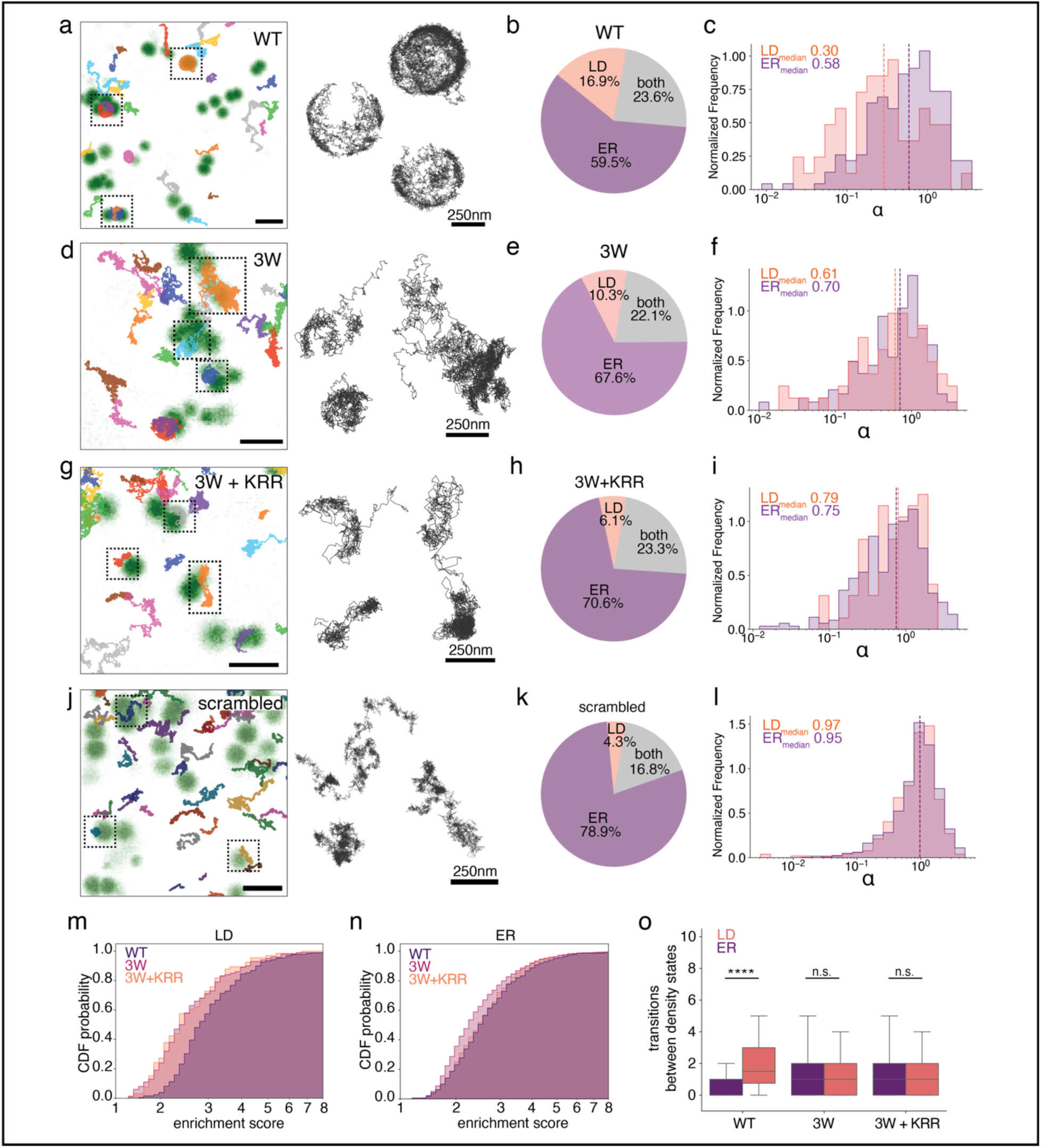
Transient interactions between specific residues and membrane nanodomains drive confinement on LDs. **a, d, g, j**. MINFLUX single-molecule trajectories for wild-type *LiveDrop* WT (a), the tryptophan mutant (3W) (d), the tryptophan and positively charged mutant (3W+KRR) (g), and the scrambled hairpin variant (j), overlaid on confocal images of LDs. Representative zoomed-in examples of LD-localized trajectories for each construct are shown on the right. Scale bars: 1 µm (left), 250 nm (right). **b, e, h, k.** Subcellular distribution of trajectories for each construct, categorized as ER-localized (purple), LD-localized (salmon), or spanning both compartments (gray). Pie charts show distribution for *LiveDrop* WT (b), 3W (e), 3W+KRR (h), and scrambled (k). **c, f, i, l.** Anomalous diffusion analysis of *LiveDrop* WT (c), 3W (f), 3W+KRR (i), and scrambled (l). The anomalous exponent (α) was calculated by fitting a power-law function to the full MSD curve of each trajectory. Distributions are shown for ER-localized (purple) and LD-localized (salmon) tracks. Dotted lines indicate medians. *N*_WT_ = 234, *N*_3W_ = 212, *N*_3W+KRR_ = 289, *N*_scrambled_ = 1584. **m, n.** Cumulative density function probabilities of spot enrichment scores were plotted for the LD (m) and ER (n) localized single molecule trajectories of *LiveDrop* variants. Enrichment scores were calculated by taking the ratio of number of single molecule localizations within each dense region and the area of the dense region. **o.** number of consecutive dense segment transitions were plotted as boxplot. Segments were labeled as dense if 70% of the total localization were within dense nanodomains. An unpaired Mann Whitney test was performed to assess statistical significance. *P* values: 0.000016 (WT), 0.47 (3W), 0.42 (3W+KRR).

Previous studies showed that the targeting of *LiveDrop* to LDs relies on tryptophan and positively charged residues^8^. To test whether these residues drive sub-diffusive behavior, we mutated the basic (*LiveDrop “*KRR”) or bulky hydrophobic (*LiveDrop “*3W”) residues and analyzed molecular motion with MINFLUX (Extended Data Fig. 3g). Mutating tryptophans did not prevent LD access nor alter D_app_ (Fig. 2d, e; Extended Fig. 3h, i), but a reduction in α upon LD localization was not found (Fig. 2f), indicating that confinement of *LiveDrop* at LDs requires these tryptophans. Consistently, the 3W mutant exhibited more dynamic transitions between LDs and the ER (Fig. 2d; Extended Data Fig. 3b). More extensive mutations of *LiveDrop*, such as combining tryptophan and basic residue mutations, essentially abolished LD targeting (Fig. 2g–i; Extended Fig. 3j,k). Instead, we found only rare LD-localized tracks in small ER-adjacent subregions. Similarly, a hairpin mutant with the same amino acids scrambled in their sequence (*LiveDrop* “scrambled”) showed similar defects and was enriched within subregions near LDs (Fig. 2j–l), indicating that both the presence and positioning of tryptophan and charged residues are essential for stable LD localization.

To assess whether specific residues mediate nanodomain interactions at LDs, we compared nanoscale localization of WT *LiveDrop* and mutant versions. WT showed higher spot and KDE density enrichment on LDs compared to the ER, indicating more spatially restricted nanodomain localization with lower α values (Fig. 2m, n; Extended Data Fig. 4a, b). Mutating tryptophans reduced density enrichment on LDs and lowered α differences between compartments, suggesting weaker, less specific nanodomain engagement (Fig. 2m, Extended Data Fig. 4a, b). Interestingly, mutating tryptophan residues led to pronounced effects at the ER where this mutant exhibited higher density enrichment and longer dwell times within ER nanodomains than WT (Extended Data Fig 4c). On the LD surface, however, these mutations led to a loss of nanodomain selectivity, resulting in reduced dynamic sampling of the nanodomains, and fewer transitions into and out of nanodomains than WT (Extended Data Fig. 4d, Fig. 2o). Since the 3W+KRR mutant does not efficiently reach LDs, we observed minimal localization enrichment (Fig. 2m, Extended Data Fig. 4d). These findings support a role for conserved tryptophans in promoting LD-specific nanodomain interactions that are reflected in the confinement of molecular motion.

To determine if specific residues mediate nanodomain interactions on LDs, we compared the nanoscale localization behavior of WT *LiveDrop* and mutant variants. WT exhibited higher spot and KDE density enrichment on LDs than the ER, consistent with more spatially restricted nanodomain localization and reduced α values (Fig. 2m, n; Extended Data Fig. 4a, b). Mutating the conserved tryptophans diminished LD enrichment and reduced α differences between compartments, indicating weaker and less selective nanodomain engagement (Fig. 2m, Extended Data Fig. 4a, b). On the LD surface, however, the mutant lost nanodomain selectivity, leading to reduced dynamic sampling and fewer transitions into and out of nanodomains (Extended Data Fig. 4d, Fig. 2o). Interestingly, these mutations had a more pronounced effect on the ER, where the 3W mutant showed elevated density enrichment and longer dwell times within ER nanodomains compared to WT (Extended Data Fig. 4c). Consistent with its ER retention, the 3W+KRR mutant failed to reach LDs and showed minimal localization enrichment (Fig. 2m, Extended Data Fig. 4d), underscoring the role of tryptophan residues in mediating LD-specific nanodomain interactions and molecular confinement.

## LD cargo targets all LDs via movement across ER-LD membrane bridges

We hypothesized that the change in motion behavior between ER and LD could be used to better understand the route of *LiveDrop* targeting to LDs, specifically by extracting the coordinates where the protein crossed the bilayer-monolayer membrane contiguity connecting the organelles. However, because these events are rare, and because MINFLUX can only track one molecule at a time, we turned to a different system.

To increase the likelihood of observing the ER to LD transition of molecules, we sought to synchronize trafficking by combining single-molecule tracking with a LD-RUSH system^17^ (Fig. 3b). We co-expressed an ER hook and *LiveDrop* fused to a streptavidin-binding peptide (SBP-Livedrop), which retained *LiveDrop* at the ER membrane even after treatment of cells with oleic acid–containing medium overnight (Fig. 3 b,c). Subsequent addition of biotin to the culture medium released SBP*-LiveDrop* from the anchor, enabling visualization of ER-to-LD protein trafficking. Within 10 min of release, *LiveDrop* relocalized from the ER to essentially all LDs in the cell, where it continued to accumulate over the next 60 mins (Extended Data Fig. 5a, b). After biotin release, SBP-*LiveDrop* localized to LDs irrespective of their size or time of formation— whether nascent or mature—indicating a uniform mechanism that facilitates and preserves membrane continuity between the ER and LDs (Extended Data Fig. 5c-e).

**Fig. 3.**
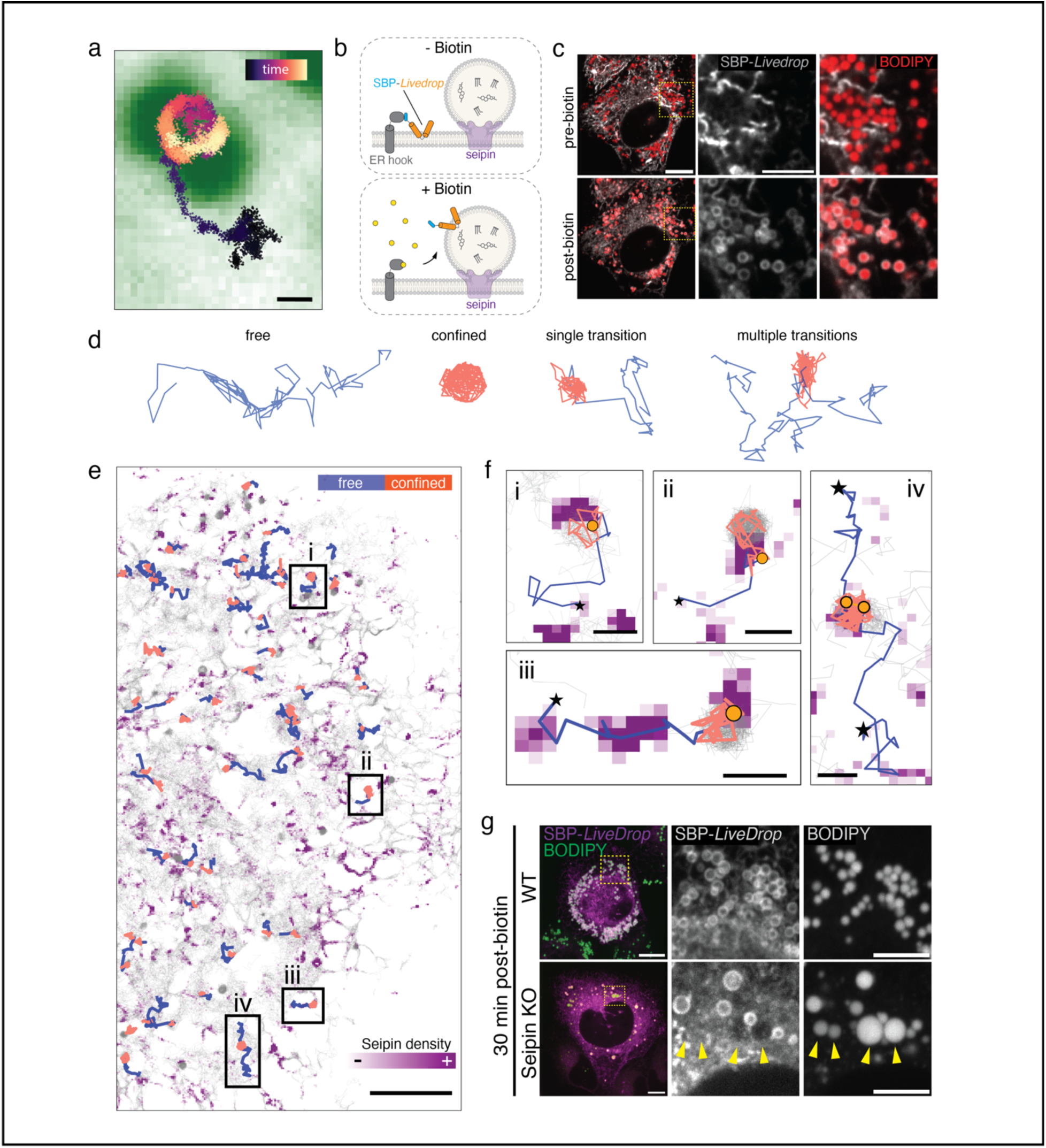
SBP-*LiveDrop* accesses LDs via seipin-mediated ER–LD contact sites. **a.** Representative MINFLUX single-molecule trajectory of SBP-*LiveDrop* entering the LD surface. Time progression is color-coded (black: start, yellow: end). LDs (green) are stained with BODIPY 493/503. Scale bar: 250 nm. **b.** Schematic of synchronized SBP-*LiveDrop* trafficking from the ER to LDs using the RUSH system. SBP-*LiveDrop* is retained on the ER membrane through interaction with an ER-localized hook. Biotin addition competes for streptavidin binding, releasing SBP-*LiveDrop* and enabling real-time monitoring of its trafficking to the LD surface. **c.** Live-cell confocal images of SBP-*LiveDrop* before and 10 min after biotin-induced release. Cells were pretreated with 250 µM oleic acid overnight to induce LD formation. SBP-LiveDrop was labeled with JFX554 Halo ligand and released by adding 80 µM biotin. LDs were stained with 1 µM BODIPY 493/503. Scale bars: 10 µm (left), 5 µm (middle). **d.** Motion types of SBP-*LiveDrop* classified by DeepSPT. Single-molecule trajectories were categorized into four groups: free-only (blue), confined-only (orange), single transition (free-to-confined or vice versa), and multiple state-switching trajectories. **e.** SBP-*LiveDrop* single-molecule trajectories plotted over a seipin density map. Cells expressing sparsely labeled SBP-*LiveDrop* and endogenously tagged seipin were imaged simultaneously using HILO microscopy. An LD prediction algorithm identified trajectories moving from the ER to LDs (see Methods). Motion states are color-coded; other tracks are shown in light grey. Scale bar: 10 µm. **f.** Examples of SBP-*LiveDrop* trajectories entering LDs through seipin-rich regions. Trajectories are color-coded by motion state (blue: free; orange: confined). Stars denote starting positions; orange circles indicate free-to-confined transitions used as proxies for LD entry. Scale bar: 500 nm. **g.** Confocal images of SBP-*LiveDrop* in wild-type and seipin knockout cells. Cells were incubated overnight with 250 µM oleic acid and imaged by spinning-disk confocal microscopy 30 min after biotin release. Arrowheads indicate LDs in seipin KO cells that failed to recruit SBP-*LiveDrop*, consistent with a loss of ER–LD membrane contact. Scale bars: 10 µm (left), 5 µm (right).

We recorded ∼56,000 SBP*-LiveDrop* single-molecule trajectories of sparsely labeled SBP-*Livedrop* by using highly inclined and laminated optical (HILO) sheet microscopy at various times after biotin release. To categorize single-molecule tracks based on their localization and anomalous diffusion behavior, we implemented DeepSPT, a machine-learning-based diffusion analysis pipeline^18^ (Fig. 3d; see Methods). In agreement with MINFLUX measurements, fitting of HILO tracks showed a lower α exponent on LDs than at the ER, indicating nano-confinement on the monolayer surface (Extended Data Fig. 6 a,b). This analysis revealed that tracks moving from the ER to LDs changed behavior from free to nano-confined motion, enabling temporal identification of ER and LD trafficking (Extended Data Fig. 6c,d).

With DeepSPT, we also captured a few instances of multiple different tracks appearing to access LDs from different directions within a short time (Extended Data Fig. 7a). To test whether one or multiple ER-LD membrane bridges allow protein trafficking, we analyzed the coordinates of LD access points where the switch from ER to LD motion pattern occurs. The motion switch points entered BODIPY-stained LD at similar locations (Extended Data Fig. 7 b,c), suggesting that each of these molecules accessed the LD through the same entry site.

## *LiveDrop* cargo protein accesses LDs via seipin-mediated membrane bridges

Seipin oligomers are components of the LD assembly complex, forming a single stable focus at ER-LD interfaces^9,12,19,20^ (Extended Data Fig. 8a). To test whether the entry point for SBP*-LiveDrop* to LDs was at seipin oligomers, we simultaneously imaged single SBP*-LiveDrop* molecules and endogenously tagged seipin after biotin release. LD-associated seipin was far less mobile than free seipin likely due to its role in maintaining the ER-LD membrane contacts^9,12,21,22^ (Extended Data Fig. 8b,c). Therefore, seipin’s single molecule positions were densely clustered at ER-LDs contact sites (Extended Data Fig. 8d,e). DeepSPT analysis of SBP*-LiveDrop* motion characteristics revealed changing motion signatures at seipin-dense regions (Fig. 3e, Extended Data Fig 9a, b; see Methods). As these tracks transitioned from the ER to LDs, their motion breakpoints colocalized at the focus formed by a stable LD-associated seipin cluster (Fig 3f), indicating protein movement through a seipin complex.

Seipin-deficiency leads to defects in LD formation and morphology and LDs detaching from the ER membrane^9,12^. Examining behavior of SBP*-LiveDrop*-RUSH in seipin-knockout cells, we found that, upon release from the ER, SBP*-LiveDrop* targeted some LDs, whereas other LDs did not receive SBP*-LiveDrop* after 30 min of biotin release, indicating a loss of bilayer-monolayer membrane bridges between the two compartments (Fig 3g). It is unclear how some LDs retain continuity with the ER in the absence of seipin; these LDs may receive *LiveDrop* via transient ER– LD contacts or by budding of small *LiveDrop*-containing LDs that subsequently fuse with mature LDs.

## Cargo movement across seipin complexes is bidirectional

Whether LD proteins can move back to the ER in human cells has been unknown. Single-molecule tracking enabled us to address this question. Since in wild-type cells, seipin maintains a membrane bridge between ER and LDs that allows proteins to cross, we tested whether these bridges allow *SBP-LiveDrop* to move back from LDs to the ER. We performed a “reverse-RUSH” experiment (Fig. 4a) in which we incubated the SBP*-LiveDrop*-RUSH-expressing cells with biotin and oleic acid overnight to stimulate LD production and allow SBP*-LiveDrop* to accumulate at LDs. To allow ER hooks to re-capture pre-labelled SBP*-LiveDrop* molecules that escaped from LDs back to the ER, we added excess avidin into the oleate-containing medium the following day to sequester biotin. After overnight avidin incubation, almost all SBP*-LiveDrop* returned to the ER (Fig. 4b). Cellular levels of SBP*-LiveDrop* did not change after avidin incubation, suggesting that the change in localization was not due to protein turnover (Extended Data Fig. 10a, b).

**Fig. 4.**
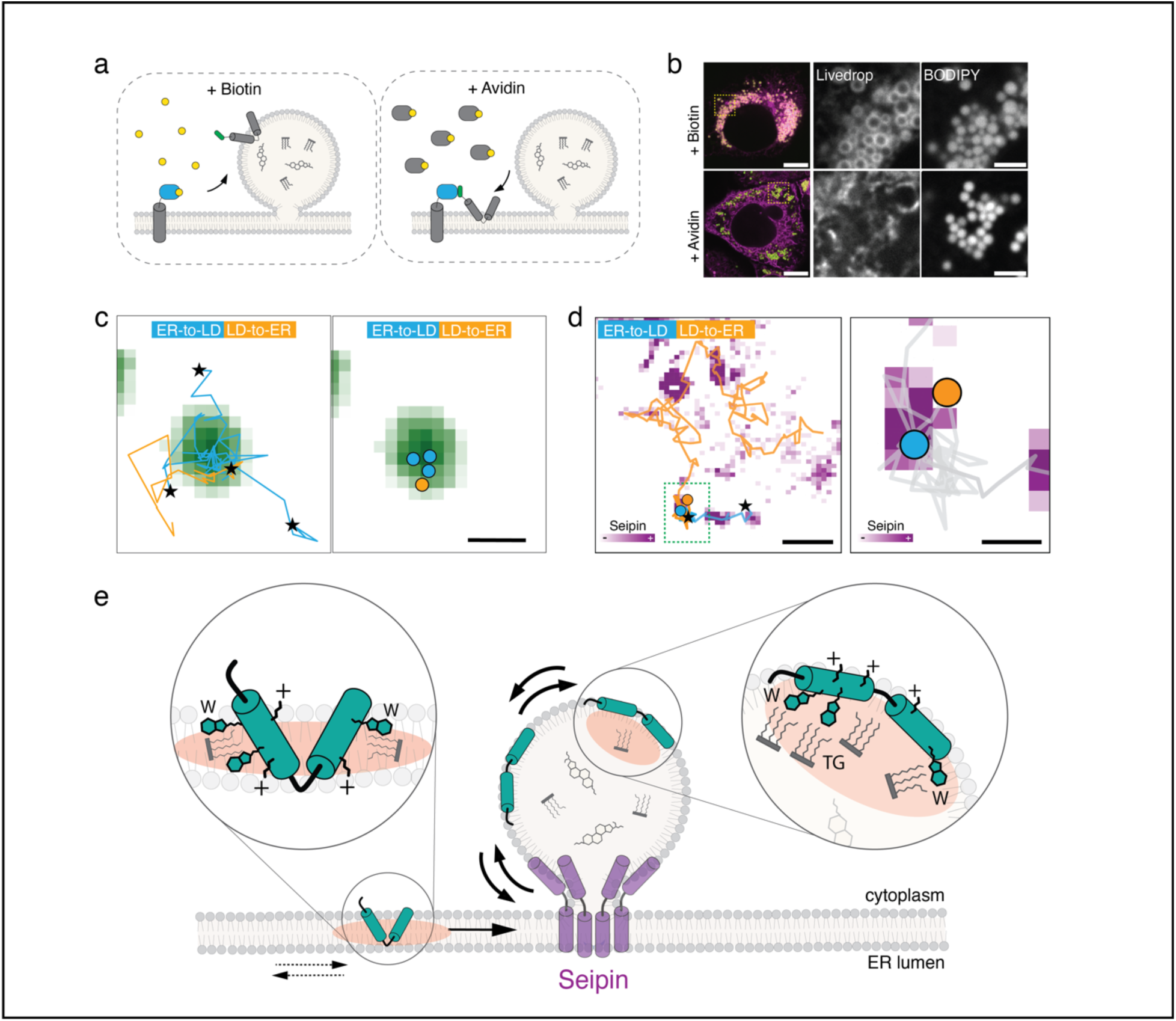
Cargo movement across seipin-mediated ER-LD contact sites is bidirectional. **a.** Schematic of the “reverse RUSH” assay. Cells were cultured in biotin-containing medium to allow SBP-*LiveDrop* accumulation on LDs. Subsequent addition of avidin displaced biotin from the ER hook, enabling re-capture of any SBP-*LiveDrop* molecules that returned to the ER membrane. **b.** Confocal images of SBP-*LiveDrop* in cells incubated overnight with biotin (top) or after avidin treatment (bottom). Cells were treated with biotin and 250 µM oleic acid overnight to induce LD formation. LD-localized SBP-*LiveDrop* was labeled with 100 nM JFX554 prior to avidin addition. After an overnight avidin incubation, cells were imaged using confocal microscopy. LDs were stained with BODIPY 493/503 (green). Scale bars: 10 µm (left), 2 µm (right). **c.** Representative MINFLUX single-molecule tracks of SBP-*LiveDrop* exhibiting bidirectional trafficking between the ER and LDs. Left: multiple trajectories showing both entry into and exit from the same LD. Right: spatial coordinates of motion switches are plotted for ER-to-LD (blue) and LD-to-ER (orange) events, overlaid on the LD channel. Stars indicate trajectory start positions. Scale bar: 500 nm. **d.** Example of SBP-*LiveDrop* bidirectional movement across a seipin-containing ER-LD contact site. Motion switch coordinates in both directions colocalize with a seipin density hotspot near the LD. Scale bars: 1 µm (left), 250 nm (right). **e.** Hairpin-containing membrane proteins exhibit comparable diffusion dynamics on both ER and LD surfaces but become progressively confined upon reaching the LD monolayer. Confinement on LDs arises from repeated, selective interactions with nanoscale domains enriched in triglycerides and phospholipids. Specifically, bulky tryptophan residues in the hairpin domain may insert into the LD core to engage triglyceride esters, whereas flanking basic residues interact with phospholipid headgroups at the surface. These cooperative interactions create an energetically favorable environment that promotes dynamic sampling of TG-rich nanodomains and gradual protein accumulation on the LD membrane. In contrast, limited TG accessibility at the ER membrane leads to more static and prolonged protein– nanodomain interactions. Since seipin bridges permit bidirectional ER–LD exchange, we propose that nanodomain-mediated confinement is a key mechanism for the sorting and stable localization of membrane proteins on LD monolayer surface.

In agreement with these results, our single-molecule tracking dataset contained instances of SBP*-LiveDrop* molecules that travelled from LD to the ER and transitioned from nano-confined to free movement (Fig. 4c, Extended Data Fig. 10c). Such LD-to-ER tracks were present at every time point measured, including ones with a short biotin incubation and later stages of biotin release (Extended Data Fig. 10d). The diffusion breakpoints of the reverse tracks were found at sites of LD entry and colocalized with seipin-dense regions (Fig. 4c, d, Extended Data Fig. 10e), indicating bidirectional movement of proteins across these regions.

## Discussion

Here we combined single-molecule analysis with unprecedented temporal and spatial resolution to address longstanding questions concerning membrane protein targeting to LDs. Our results provide a mechanistic framework in which ER-resident membrane proteins are not simply trafficked to LDs but are selectively retained through nanoscale interactions at the LD surface. The observations that GPAT4, HSD17B13, and *LiveDrop* diffuse with similar speeds in both the ER and LD membranes yet become confined on LDs suggest that the LD monolayer imposes spatial constraints that promote gradual protein accumulation. Consistent with this, we observed that *LiveDrop* moves bidirectionally across seipin-containing ER–LD bridges, supporting that protein accumulation is driven by local retention rather than physical barriers. This mechanism allows for dynamic and selective enrichment of key enzymes on LDs, with implications for metabolic regulation and organelle specialization. Such a mechanism represents an efficient strategy for inter-organelle communication, particularly in systems that lack dedicated protein translocation machinery.

The mechanism underlying the interaction of LD proteins with nanodomains is currently uncertain. Our data suggest that tryptophans in the hairpin domain do not simply mediate binding, but fine-tune compartment-specific nanodomain interactions. The limited access to TG within the ER bilayer may create an energetic penalty that promotes exit toward the LD monolayer, where TG is abundant and accessible. Once on LDs, tryptophan-TG interactions facilitate transient, repetitive engagement with nanodomains, enabling dynamic sampling and gradual accumulation. In contrast, mutants lacking tryptophans lose this selectivity, resulting in increased ER retention and prolonged, less dynamic interactions with nanodomains (**Fig 4e**). These findings suggest that nanodomain engagement is likely driven by biophysical properties that create local heterogeneity within membranes^23,24^. Tryptophans, thus, dynamically couple molecular motion with lipid composition, enhancing compartmental specificity and driving accumulation on the LD surface. Molecular dynamics simulations further support this model, showing that central tryptophan residues (e.g., W172 in *Drosophila* GPAT4) can interact with the ester groups of TGs at LD surfaces^8^.

This mechanism of LD cargo accumulation parallels principles of protein sorting in the secretory pathway, where weak and yet specific interactions between proteins and specific lipids, such as sphingolipids and cholesterol, promote clustering of nanodomains targeted to apical membranes in polarized cells^25–28^. These findings support a model in which lipid-protein nanoclustering functions as a mechanism for selective protein sorting and targeting between cellular components.

## Acknowledgements

We thank members of the Harper and the Farese & Walther laboratories for helpful discussions, specifically Dr. Henning Arlt and Dr. Jeeyun Chung for discussions on seipin and relevant reagents, Dr. Pedro Manuel Carpio Malia for discussions on *LiveDrop* and GPAT4, and Dr. Kelsey Hickey for scientific input and helpful discussions. We thank Dr. Michelle Ocana at the Neurobiology Imaging Facility (NIF) at Harvard Medical School for MINFLUX instrumentation, Dr. Otto Wirth for MINFLUX data analysis discussions, Dr. Talley Lambert at the Center of Imaging Technologies and Education (CITE) at Harvard Medical School for confocal and HILO imaging help, and Dr. Luke Lavis for Janelia Fluor dyes. This work was supported by NIH R01 grants GM124348 (R.V.F), GM097194 (T.C.W), and NS083524 (J.W.H.), NNF challenge grant NNF23OC0081287 (NSH), NNF infrastructure grant NNF22OC0075851 (NSH), Vellux foundation 18333 (NSH) and by generous support from Ned Goodnow (J.W.H.). A.M. is a Helen Hay Whitney Foundation fellow and T.C.W. is a Howard Hughes Medical Institute investigator.

## Author contributions

A.M., T.C.W., and R.V.F. conceived, designed, and interpreted the experiments and wrote the manuscript, with input from all authors. A.M. conceptualized, designed and conducted experiments, many of which were performed during an extended stay in the laboratory of J.W.H. J.M. performed MINFLUX data acquisition for *LiveDrop* variants. A.M. performed MINFLUX data analysis. J.K.H. performed DeepSPT analysis, instance segmentation, and track classification. A.M. and J.K.H. performed image analysis, statistics, and data visualization. J.W.H. and N.S.H. provided intellectual feedback and engaged in discussions throughout.

## Data and Code Availability

All raw imaging data for confocal, HILO and MINFLUX experiments can be found at https://data.mendeley.com/preview/zbtvr59j2p?a=e9c075b7-5ef8-4c82-b1d0-f6393fd2f9df. Code and data analysis for MINFLUX tracking datasets can be found at https://github.com/amizrak/MINFLUX_analysis.git.

## Methods

### Chemicals

Janelia Fluor dyes with HaloTag JFX544 and JFX650 ^29^ were generous gifts from Luke Lavis (Janelia Research Campus). BODIPY 493/503 (D3922) and HCS Lipid TOX Deep Red Neutral Lipid Stain (H34477) were purchased from Thermo Fisher Scientific. AUTOdot (SM1000b) was purchased from Abgent. Avidin (A9275) and biotin (B4501), fatty acid-free BSA (A6003) and oleic acid (O1008) were purchased from Sigma-Aldrich.

An oleic acid stock solution was prepared at a stock concentration of 10 mM in 3 mM fatty acid–free BSA-PBS solution. The solution was incubated in 37°C shaking incubator for at least 1 h to dissolve fatty acids, filtered through a 0.22-µm filter, aliquoted and stored in −20°C.

For RUSH experiments, a 10 µM avidin stock solution was prepared by dissolving the lyophilized powder in sterile PBS and stored in −80°C. An 80 mM biotin solution (1000X) was prepared in DMSO and stored in −20°C.

### Plasmid construction

All PCRs were performed using Q5 High Fidelity DNA Polymerase (M0491-NEB) and restriction enzymes from New England Biolabs. Full-length GPAT4, HSD17B13, SBP-*LiveDrop* WT, 3W, 3W+KRR and scrambled constructs were cloned using GeneBlocks (IDT) into lentiviral backbone under control of the EF1a promoter. RUSH constructs were generated by amplifying the ER hook sequence from Str-ii vector (65300 - Addgene) and the SBP-*LiveDrop* RUSH construct from a geneBlock (IDT). The inserts were cloned into a pLVX lentiviral expression vector (134665 Addgene) using HiFi Assembly (E5520 - NEB).

### Lentiviral production

Lentivirus was produced by transfecting HEK293T with pLVX lentiviral plasmid containing RUSH constructs and standard packaging vectors using *Trans*IT-LTI Transfection Reagent (Mirus, MIR 2306). Viral particles in the supernatant were harvested 2 days post-transfection and stored at −80°C.

### Cell culture

SUM159 human breast cancer cells were a kind gift by the laboratory of Tomas Kirchhausen (Harvard Medical School) and maintained in DMEM/F-12 GlutaMAX (Life Technologies, #10565042) with 5 μg/ml insulin (Cell Applications), 1 μg/ml hydrocortisone (Sigma), 5% FBS (Life Technologies 10082147; Thermo Fisher), 50 μg/ml streptomycin, and 50 U/ml penicillin. Cells were maintained in a 5% CO_2_ incubator at 37°C and at <70% confluent. To induce TG synthesis, cells were treated with growth medium containing 250 μM oleic acid complexed with fatty acid–free BSA.

### Stable cell line generation

SUM159 cell lines stably expressing RUSH constructs were generated by lentiviral transduction. In brief, cells were seeded into six-well plates in growth medium 2 days before virus introduction. On the day of transduction, cells were treated with 8 µg/ml Polybrene (TR-1003) for 5 mins and incubated with viral supernatant for 2 days. Cells were given 4 days to express the constructs, followed by population sorting using fluorescence-activated cell sorting (FACS) on a SONY SH800S sorter. The presence of tags in sorted populations was confirmed by fluorescent microscopy prior to experiments.

### Live cell imaging

SUM159 cells were plated on 35-mm glass-bottom dishes (MatTek Corp). Before imaging, growth medium was replaced with phenol-free DMEM/F12 medium containing 15 mM HEPES, pH 7.5, 5 μg/ml insulin, 1 μg/ml hydrocortisone, 5% FBS, 50 μg/ml streptomycin, and 50 U/ml penicillin to prevent background fluorescence. For Halo tag labelling, cells were incubated with growth medium containing 100 nM Halo ligand as indicated, followed by three washes with label-free growth medium. For LD staining, cells were treated with BODIPY 493/503 (D3922, Thermo Fisher Scientific) or HCS LipidTOX™ Deep Red Neutral Lipid Stain (H34477, Thermo Fisher Scientific) at a 1:1000 dilution for 1 h prior to imaging. As indicated, 1 μg/ml Hoechst 33342 (H3570, Thermo Fisher Scientific) was added to growth medium for labelling nuclei.

Live cell imaging was performed using a Nikon Eclipse Ti inverted microscope equipped with Perfect Focus, a CSU-X1 spinning disk confocal head (Yokogawa), Zyla 4.2 Plus (Andor) or Orca-FUSION (Hamamatsu) sCMOS cameras, and controlled by NIS-Elements software (Nikon). The microscope is equipped with a live chamber (OkoLabs) to maintained cells at 37°C and 5% CO_2_. Images were acquired through 100× Plan Apo 1.40 NA objective (Nikon). Cells were excited with solid state lasers (405-, 488-, 560-, or 640-nm - Agilent). The channels of multicolor images were acquired sequentially.

### MINFLUX live cell single-molecule tracking

For single-molecule experiments, SUM159 cells that stably express LD constructs were prepared similar to above with the following modifications. To achieve single-molecule labeling, cells were incubated with 50–80 pM JFX650 ^29^ for 30 min and washed three times with media. Cells were incubated with growth medium containing 250 µM oleic acid for 3 h to induce LD formation. LDs were labelled with 1 µM BODIPY 493/503 30 min prior to imaging.

MINFLUX data were recorded on a commercial Abberior Instruments MINFLUX setup (Abberior Instruments GmbH), similar to the one reported by Schmidt et al. ^15^. The system was equipped with a 100x/1.4 NA magnification oil immersion lens, a 640-nm continuous-wave laser for exciting JFX650-labeled *LiveDrop* in both confocal and MINFLUX mode, and a 488-nm pulsed laser to image lipid droplets in confocal mode, with a pixel size of 50 nm. MINFLUX tracking was performed with the standard 2D tracking sequence provided by Abberior Instruments by increasing 640-nm laser power over the subsequent iterations. The single molecule emission was detected at 653–750 nm with the pinhole set to 0.83 AU. Before and after the actual MINFLUX tracking measurements, confocal images of the LDs were acquired to serve as a reference for the subcellular context. The Abberior Instruments Inspector software with MINFLUX drivers was used to operate the system.

### MINFLUX analysis and data visualization

MINFLUX single-molecule localization data were acquired across multiple experimental sessions and processed to extract track coordinates and assign localizations relative to LD structures using a custom python script. Raw .npy files containing X, Y, and time values for each track were extracted in Pandas. Spatial coordinates were scaled to µm using instrument-specific pixel length (50 nm) and offset parameters retrieved from metadata or auxiliary parameter files. To ensure track fidelity, tracks were filtered to exclude those with fewer than 500 time points, limited spatial dispersion, or minimal net displacement. The initial localization of each track was excluded to correct for known positional offsets at track initiation. A secondary filtering step removed tracks of which total displacement is smaller than 100 nm in both axis for the entire track as these likely represent background dye aggregates.

Confocal images of LDs were processed to generate binary masks by applying an intensity thresholding, followed by morphological dilation (5 pixels) to expand detected regions using OpenCV. Pixel coordinates corresponding to the masked LD areas were extracted, and single-molecule track positions were rescaled and compared to this mask to classify localizations as occurring inside or outside LD masks. Tracks were then grouped into three categories: those entirely inside, entirely outside, or exhibiting multiple transitions across the LD boundary. Tracks were further analyzed to compute stepwise displacements, localization precision, and time intervals between observations. Spatial and temporal characteristics of each track were visualized by overlaying trajectories on LD images and constructing density maps. All processed tracks across replicates were aggregated into a final dataset for downstream analysis. Localization precision was calculated by analyzing the standard deviation of the changes between each consecutive localization in both axes.

### Apparent and anomalous diffusion analysis of MINFLUX single molecule tracks

To capture local variations in diffusivity, a sliding window analysis was performed to calculate the mean squared displacement (MSD) and apparent diffusion coefficient (D) across individual tracks. A window size corresponding to a 5 ms time frame was determined based on the median inter-frame interval and applied to each track using a rolling approach. For each window, time lags and squared displacements relative to the initial position were computed. MSD values were derived incrementally for each lag, and the apparent diffusion coefficient was estimated from the slope of the MSD curve using the Brownian relation D=MSD/(4t). Windows with fewer than two valid lag times were excluded from analysis. The resulting time-resolved diffusion coefficients were annotated with the corresponding spatial coordinates and localization category (inside or outside LDs), enabling dynamic assessment of protein mobility across subcellular compartments.

To characterize diffusion behavior and anomalous transport, MSD curves were fit to a power-law model of the form MSD(t)=4Dt^α^. Data points within entire duration of individual track (up to 5 s) were fit using nonlinear least-squares regression in SciPy package. Fitted parameters were retained only if they satisfied quality control criteria, including physical bounds on α (0 < value < 5). Tracks were then stratified based on localization (inside vs. outside LDs), and summary statistics, including means, medians, and standard deviations and α, were computed. Distributions of fit parameters were visualized using log-scaled histograms, with kernel density overlays for the anomalous exponent, allowing comparison of diffusion dynamics between subcellular regions. Fit parameters and statistical significance were calculated using Mann-Whitney test provided by SciPy module.

### Analysis of local density dynamics

To quantify nanoscale clustering behavior of SBP-*LiveDrop*, local molecular density was analyzed across 100-ms non-overlapping segments of ER- and LD-localized single-molecule trajectories. For each segment, a two-dimensional Gaussian kernel density estimate (KDE) was computed from the (x, y) coordinates of localizations. KDE values were evaluated on a regular grid (60 pixels/µm) centered on the segment’s spatial extent with a ±0.2 µm margin. From each KDE map, the maximum, mean, and total density values (µm⁻²) were extracted. Protein-dense nanoregions were defined by applying a threshold equal to the median of all maximum KDE values across the dataset, and binary masks of dense areas were generated accordingly.

Localizations within each segment were classified as inside or outside protein-dense regions based on their KDE values. Binary traces of dense state occupancy were used to calculate the fraction of localizations in dense zones and the number of transitions between dense and non-dense states. Dense and total areas per segment were measured from the KDE masks using median threshold values, and the clustering ratio (dense/total area) as well as the spot density per µm² were computed. All features were aggregated at the segment and track levels for subsequent comparison between ER- and LD-localized trajectories. To quantify protein movement between nanodomain-associated and diffuse membrane regions, KDE segments were classified as nanodomain-associated if the ratio of localized spots exceeded 70%. Transitions between states were then computed for each track by counting the number of state changes across consecutive segments.

### LD-RUSH assay

To retain hairpin constructs at the ER membrane, cells were incubated with media containing 250 µM oleic acid and 50 nM avidin (21121, Sigma) overnight to induce LD formation and sequester free biotin in the growth medium, respectively. The next day, cells were labelled with the indicated Halo ligands, washed three times with fresh medium containing 50 nM avidin, and incubated in media containing 250 µM oleic acid, 1 µM BODIPY 493/503 and 50 nM avidin. Release from the ER was initiated by adding 80 µM of biotin (B4501, Sigma) into the imaging dish dropwise with a tubing attached to a syringe to prevent any shift in the imaging plane. Image acquisition was started immediately upon biotin release. For kinetics experiments, images were taken every 2 min unless noted otherwise.

Pulse-chase reverse RUSH experiments were performed with cells grown in medium containing 250 µM oleic acid and 1 µM of biotin overnight to allow *SBP-LiveDrop* to localize to LDs. Next day, cells were labelled with 100 nM JFX554 Halo ligand for 1 h, washed three times with fresh medium, and incubated with oleic acid–containing medium supplemented with PBS or avidin overnight to sequester any remaining biotin and allow ER hook to capture pre-labelled *SBP-LiveDrop* at the ER. At 1 h before imaging, 1 µM BODIPY 493/503 was added to cells to label LDs, and live cells were imaged with a confocal microscope.

### HILO live-cell single-molecule tracking

Single-molecule imaging was performed using Nikon Ti motorized inverted microscope equipped with Agilent MLC 400B Laser unit (405/488/561/635), ImageEM EM-CCD camera (Hamamatsu) fitted with a 100×, NA 1.49 Apo TIRF objective lens (Nikon) combined with a 1.5x tube lens (image pixel size 0.107 μm). The angle of illumination was optimized to achieve full depth-of-focus of the objective lens with low SNR. Simultaneous dual color imaging experiments were performed with a dual view mirror setup (Optical Insights, AZ) equipped with either a 565LP dichroic mirror in series with 525/50 (left) and 600/50 (right) (for JFX554) or a 610LP dichroic mirror followed by 525/50 (left) and 645/75 (right) (for JFX650) Chroma filters for optimal channel separation. Single-molecule images were captured with EM gain at 200 and contrast gain at 2 with 30-ms exposure time combined with no delay acquisition to achieve a framerate of 31 fps.

### HILO Image quantification and track generation

All acquired images were processed and prepared for figures using Fiji ^30^. Confocal images were quantified using CellProfiler ^31,32^software with a custom pipeline, including image enhancement, object identification for LD, mask segmentation and quantification of channel intensities to measure SBP-*LiveDrop* intensity on LDs.

Single-molecule tracks were generated using the Fiji TrackMate plugin ^33,34^. Tracks were generated using LAP tracker with a 0.5-µm search radius for each single-molecule spot combined with gap filling of four frames within 0.7-µm radius. Initial filtering was performed using a custom Python script to filter out tracks shorter than 10 frames. Combined tracks were exported in .csv format and analyzed in DeepSPT modules (below).

### Instance segmentation images to capture LDs

To accurately segment LDs with varying sizes, we utilized a deep-learning framework Stardist^35^, which is pre-trained to be a versatile segmentation tool in fluorescent microscopy. In each frame, identified LDs were converted into a binary mask of zeros for background and a single unique integer for the pixels it occupies. Only LDs of fewer than 150 pixels were considered unless LDs were inseparably close or seipin has been knocked out, promoting LDs, in which case large masks were kept. The unique masks identified per frame were tracked in time using Trackpy’s nearest neighbor linking, with a search range of 5 or 10 pixels depending on visual evaluation and, thus, resulting in a time-persistent unique identifier per LD. The annotation of pixels to either ER or a time-persistent unique LD identifier enables the evaluation of the cellular localization of each SBP-*LiveDrop* track with individual LD resolution.

### Deep-learning-assisted analysis of temporal diffusional behavior

To characterize the heterogeneous motion of SBP-*LiveDrop*, DeepSPT, a modular deep learning framework specifically designed for interpreting heterogeneous single-particle trajectories, was employed. DeepSPT consists of three core modules: (1) temporal segmentation of diffusional behavior, (2) extraction of trajectory-level diffusion features, and (3) a downstream classifier trained on experimental data to perform system-specific classification tasks.

### Temporal segmentation and quantification of diffusional behavior

SBP-*LiveDrop* trajectories were input directly into DeepSPT’s temporal segmentation module, which predicts a diffusional state and associated confidence score at each time point. Diffusion modes observed on ER and LD membranes are diverse, but generally more restricted on LDs due to their limited area. To simplify behavioral classification, diffusion states were binned as either *“Free”* (Brownian or directed motion) or *“Restricted”* (subdiffusive or confined motion). These frame-wise predictions were used to identify diffusion state transitions and evaluate how these dynamics correlated with cellular localization. In parallel, trajectory-level features were extracted using DeepSPT’s second module, including: diffusion coefficient (estimated from single time interval MSD), anomalous diffusion exponent (α, from MSD = 4Dt^α^), explored area, and frequency of free versus restricted states.

### DeepSPT feature representation of SBP-LiveDrop trajectories

Features derived from the first two modules were concatenated into fixed-length trajectory embeddings that included: the number and sequence of transitions between free and restricted states, global diffusion metrics, and averaged features within each state. These representations were then used to train DeepSPT’s downstream classifier to distinguish ER vs. LD-localized tracks. The classifier consisted of an ensemble of three Random Forest models, each with 100 trees and maximum depths of 5, 6, or 7. Final predictions were made by majority vote. To mitigate class imbalance—due to most SBP-*LiveDrop* tracks residing in the ER—random oversampling of LD-classified tracks was performed. Classifier performance was evaluated using leave-one-group-out cross-validation, where each biological replicate (i.e., independent HILO experiment) was treated as a separate group, ensuring the model generalized across experimental conditions and cell-to-cell variability.

### Classification of trajectory cellular location solely by diffusion and model validation

DeepSPT’s third module utilizes the feature representation derived from the first two modules to provide classification directly using experimental data. The downstream classifier employed for the prediction of SBP-*LiveDrop’s* cellular localization consists of an ensemble of three Random Forest classifiers each consisting of 100 tree estimators and with maximum tree depth of 5, 6, and 7, respectively. The final ensemble prediction is taken as the mode prediction of the three classifiers. To overcome biased predictions due to class imbalance as most SBP-*LiveDrop* tracks are on the ER, random oversampling of the minority class is implemented. Model performance was evaluated using leave-one-group-out cross-validation where each group is a biological replicate (i.e., an independent TIRF single-particle-tracking experiment) to confirm the model’s ability to generalize across experimental variability and to unseen cells.

### Identification of LD-associated seipin

To identify LD-associated seipin assemblies, we combined diffusional behavior classification with spatial density mapping. Tracks classified as >90% free-diffusing by DeepSPT were excluded. Remaining tracks were converted into a 2D positional histogram to generate a seipin density heatmap. This heatmap was then binarized using a dataset-specific threshold and processed with Laplacian of Gaussian filtering (via scikit-image^36^), using a maximum sigma of 2 and a detection threshold of 0.1, to identify discrete regions of high-density, presumed to be LD-associated seipin foci.

### Evaluation of linking error potential

Evaluation of linking error potential was performed by calculating the distance from every individual SBP-*LiveDrop* detection in frame, *i*, to every detection in frame, *i+1*, and counting the number of detections within the SBP-*LiveDrop* tracking search range.

### Statistics

Unless otherwise stated, results are presented as mean ± standard deviation. Statistical analyses of results were performed using SciPy statistics package in Python. Statistical tests used for analysis are indicated in figure legends. Statistically significant calculations were denoted with * for p<0.05, ** for p<0.01, *** for p<0.001, **** for p<0.0001. Actual *P* values are included in the figure legends.

---

**Extended Data Fig. 1.**
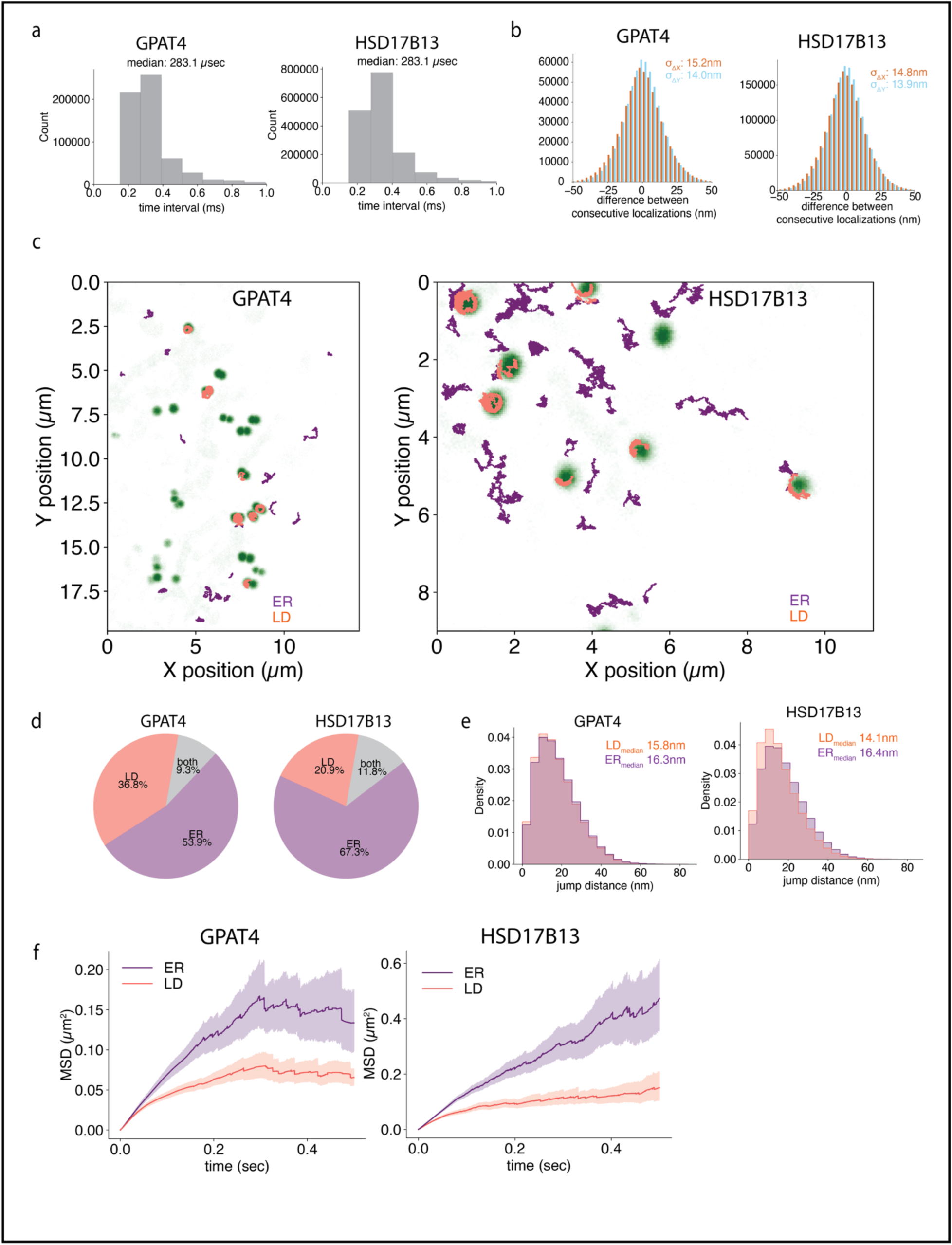
MINFLUX single-molecule analysis of metabolic enzymes GPAT4 and HSD17B13 on the ER and LDs. **a.** The sampling rate of MINFLUX was determined by calculating the time interval between consecutive localizations of individual GPAT4 and HSD17B13 molecules. **b.** The difference between consecutive localizations within single-molecule MINFLUX tracks for GPAT4 (left) and HSD17B13 (right). Standard deviation along each axis was calculated as a proxy for localization precision. **c.** Representative image showing classification of single-molecule tracks based on subcellular localization. Molecules localized within the LD mask were classified as LD-associated (orange), and those outside the mask were assigned to the ER (purple). **d.** Proportion of single-molecule tracks for GPAT4 (left) and HSD17B13 (right) localized to LDs (orange), ER (purple), or spanning both compartments (gray). **e.** Distribution of single-molecule jump distances for GPAT4 (left) and HSD17B13 (right) on the ER (purple) and LD surface (orange). Median values are indicated. **f.** Mean-square displacement (MSD) curves of individual ER- and LD-localized tracks for GPAT4 (left) and HSD17B13 (right), plotted as a function of time interval.

**Extended Data Fig. 2.**
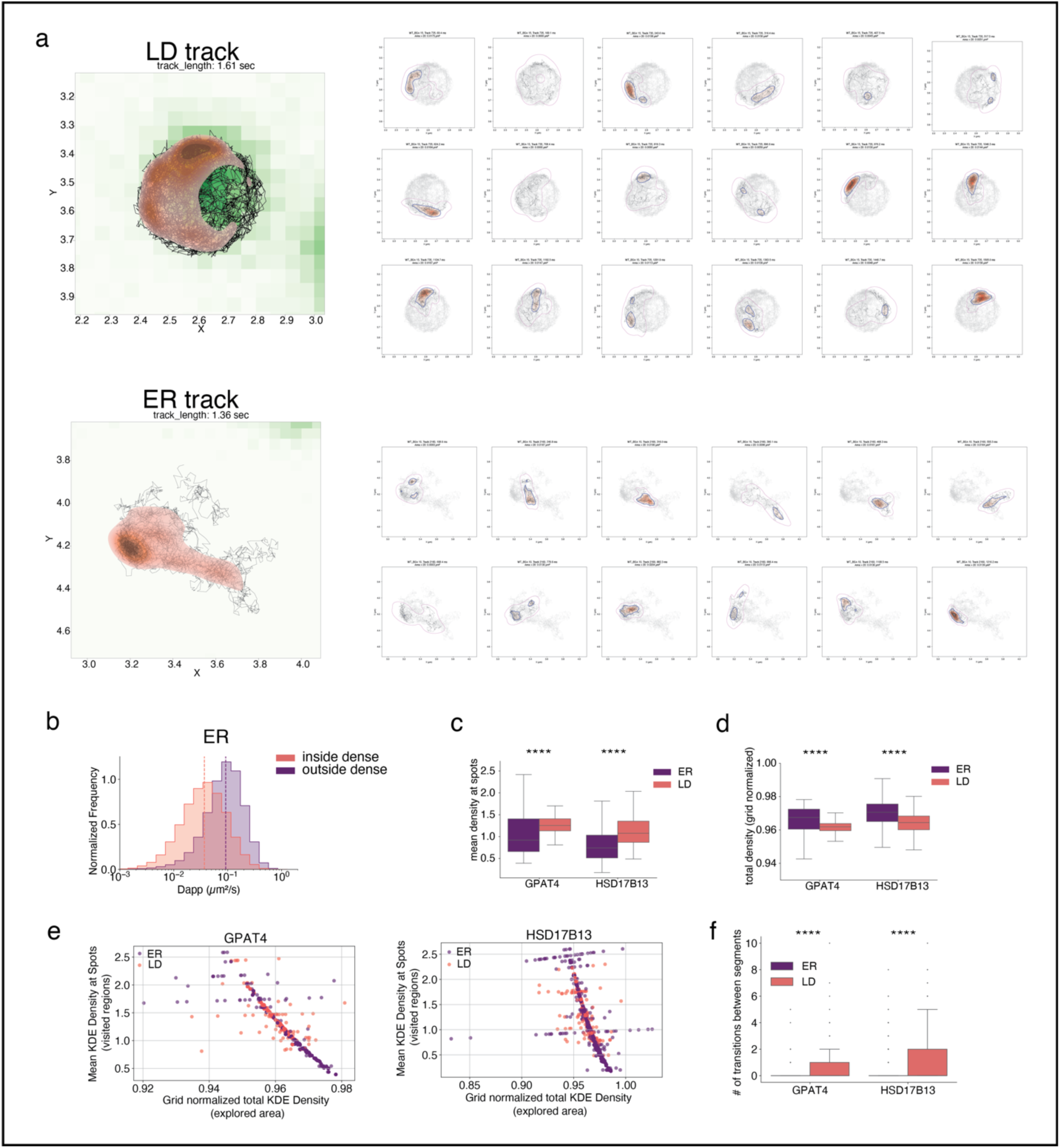
Nanodomain association of model LD cargo analyzed using Gaussian KDE. **a.** Representative examples of GPAT4 tracks localized to LDs (top) and the ER (bottom), showing full-track KDE density maps (left). Tracks were segmented into 100 ms windows, and regions exceeding the dataset’s median KDE density were defined as “dense.” The area of these dense regions was quantified. **b.** Apparent diffusion coefficients were calculated for ER-localized tracks using a 5-ms rolling window and clustered by whether segments were inside (orange) or outside (purple) dense regions. Median values are indicated by dotted lines. **c.** Mean KDE density within dense regions was computed by dividing the summed KDE values within dense regions by the number of localizations per segment, for ER (purple) and LD (orange) tracks. Unpaired t-tests revealed significant differences: *P* = 1.15 × 10⁻¹⁸ (GPAT4), 1.88 × 10⁻⁸³ (HSD17B13). **d.** Total KDE density per segment was normalized to the grid area and compared between ER (purple) and LD (orange) tracks. Unpaired t-tests: *P* = 1.57 × 10⁻¹⁷ (GPAT4), 1.21 × 10⁻⁶⁵ (HSD17B13). **e.** Density enrichment, calculated as the ratio of mean KDE density within dense regions to total normalized KDE density per segment, is shown for ER (purple) and LD (orange) tracks. **f.** The number of transitions between dense and non-dense regions per 100-ms segment was quantified for GPAT4 and HSD17B13 and plotted as boxplots for ER (purple) and LD (orange) localizations. Unpaired t-tests: *P* = 0.00013 (GPAT4), 9.4 × 10⁻⁹ (HSD17B13).

**Extended Data Fig. 3.**
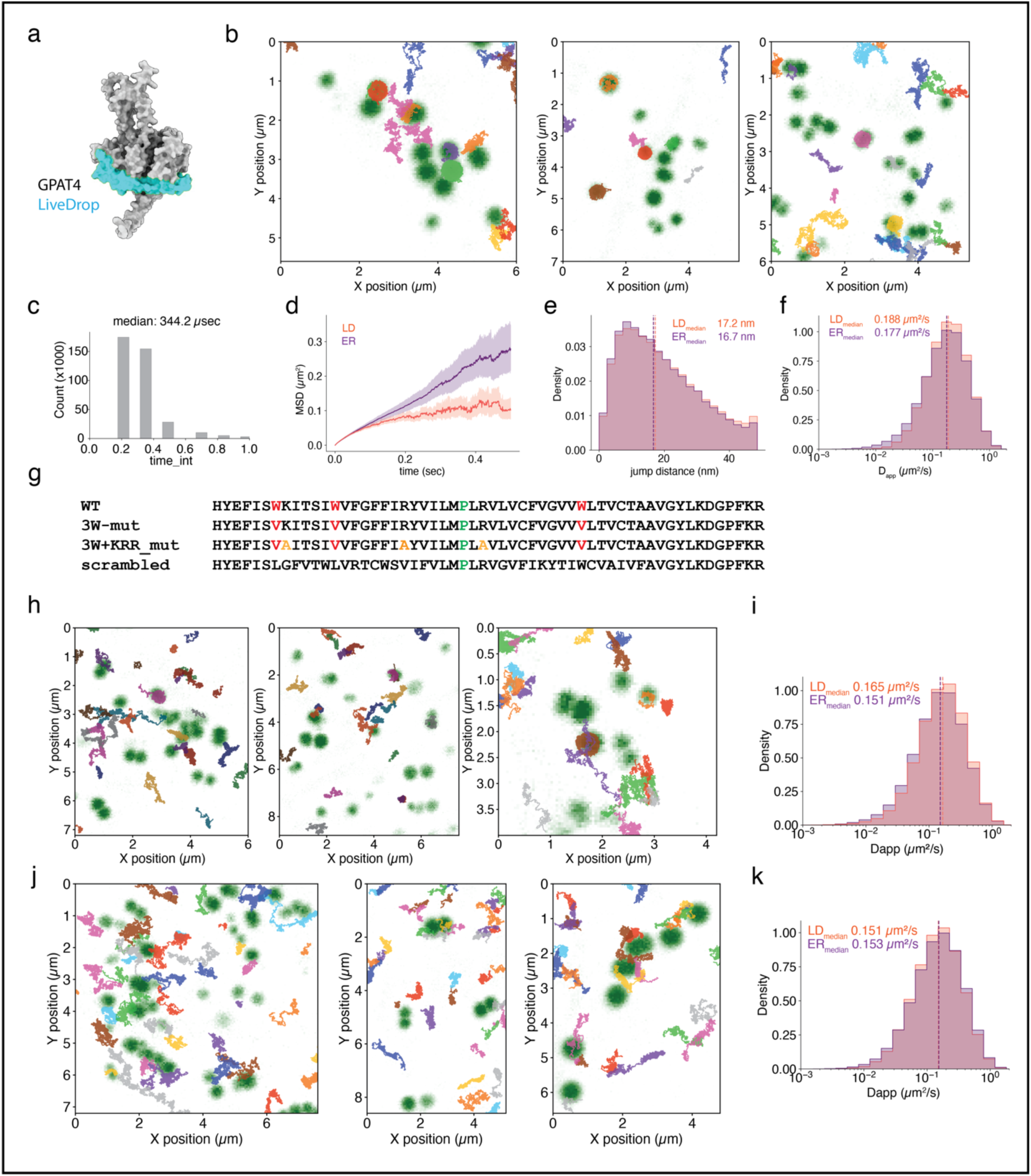
Molecular motion analysis of *LiveDrop* and mutant hairpins using MINFLUX nanoscopy. **a.** AlphaFold-predicted structure of GPAT4, with the engineered hairpin region (“*LiveDrop*”) highlighted in blue. **b.** Representative MINFLUX single-molecule tracks of *LiveDrop* in human cells, with individual trajectories shown in different colors. **c.** Sampling rate distribution of MINFLUX tracking for *LiveDrop*. **d.** Mean squared displacement (MSD) curves of *LiveDrop* single-molecule tracks, colored by subcellular location: ER (purple) and LD (orange). **e.** Jump distance distributions for *LiveDrop* tracks localized to the ER (purple) or LD (orange). Median values are indicated by dotted lines. **f.** Apparent diffusion coefficients were calculated using a 5-ms rolling window for ER-(purple) and LD-localized (orange) *LiveDrop* tracks. **g.** Hairpin sequences of *LiveDrop* and its mutants used in this study. The hinge region (green) was preserved to maintain hairpin integrity. **h.** Representative MINFLUX tracks of the 3W mutant, with each trajectory colored individually. **i.** Apparent diffusion coefficients of 3W mutant tracks calculated with a 5-ms rolling window; medians are indicated by dotted lines. **j.** Representative MINFLUX tracks of the 3W+KRR mutant, with individual trajectories colored. **k.** Apparent diffusion coefficients of 3W+KRR mutant tracks using a 5-ms rolling window. Median values are indicated by dotted lines.

**Extended Data Fig. 4.**
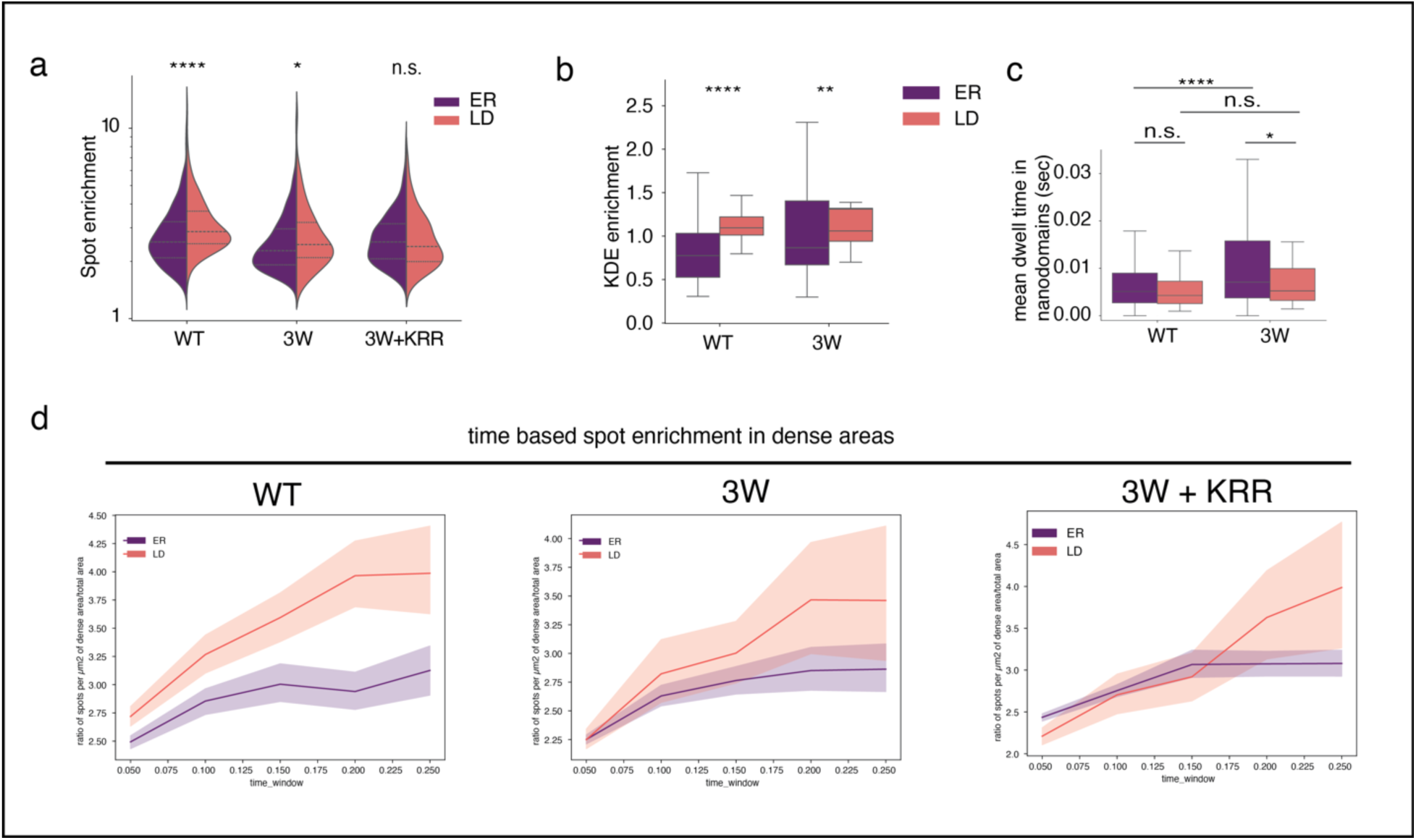
Specific amino acid sequences in the hairpin region mediate nanodomain interaction. **a.** Spot enrichment within dense membrane nanodomains was quantified by calculating the ratio of the number of localizations per unit area in dense regions to that in the total KDE area. This enrichment ratio was plotted as split violin plots comparing ER-localized (purple) and LD-localized (orange) trajectory segments for wild-type (WT) and mutant hairpins. Differences in distribution were assessed using an unpaired t-test. *P* values: 2.9×10^−8^ (WT), 0.032 (3W), 0.88 (3W+KRR). **b.** KDE-based density enrichment within nanodomains was quantified for WT LiveDrop and the 3W mutant. Enrichment was defined as the ratio of the mean KDE value at each dense region to the mean KDE of the full grid. Data were plotted as box plots for ER and LD membranes. *P* values: 9.11×10^−21^ (*LiveDrop*), 0.004 (3W). **c.** Average dwell times inside nanodomains for WT and 3W mutant were plotted as a box plot. The ER (purple) and LD (orange) localized tracks within each dense segment was calculated based on time spent in each region. *P* values: 1×10^−5^ (WT_ER_ vs 3W_ER_), 0.065(WT_LD_ vs 3W_LD_), 0.17 (WT_ER_ vs WT_LD_), 0.049 (3W_ER_ vs 3W_LD_). **d.** Time-resolved density enrichment was plotted for WT, 3W, and 3W+KRR trajectories. Segment lengths were varied to assess temporal trends, and enrichment was calculated as described in (a) for each time window.

**Extended Data Fig. 5.**
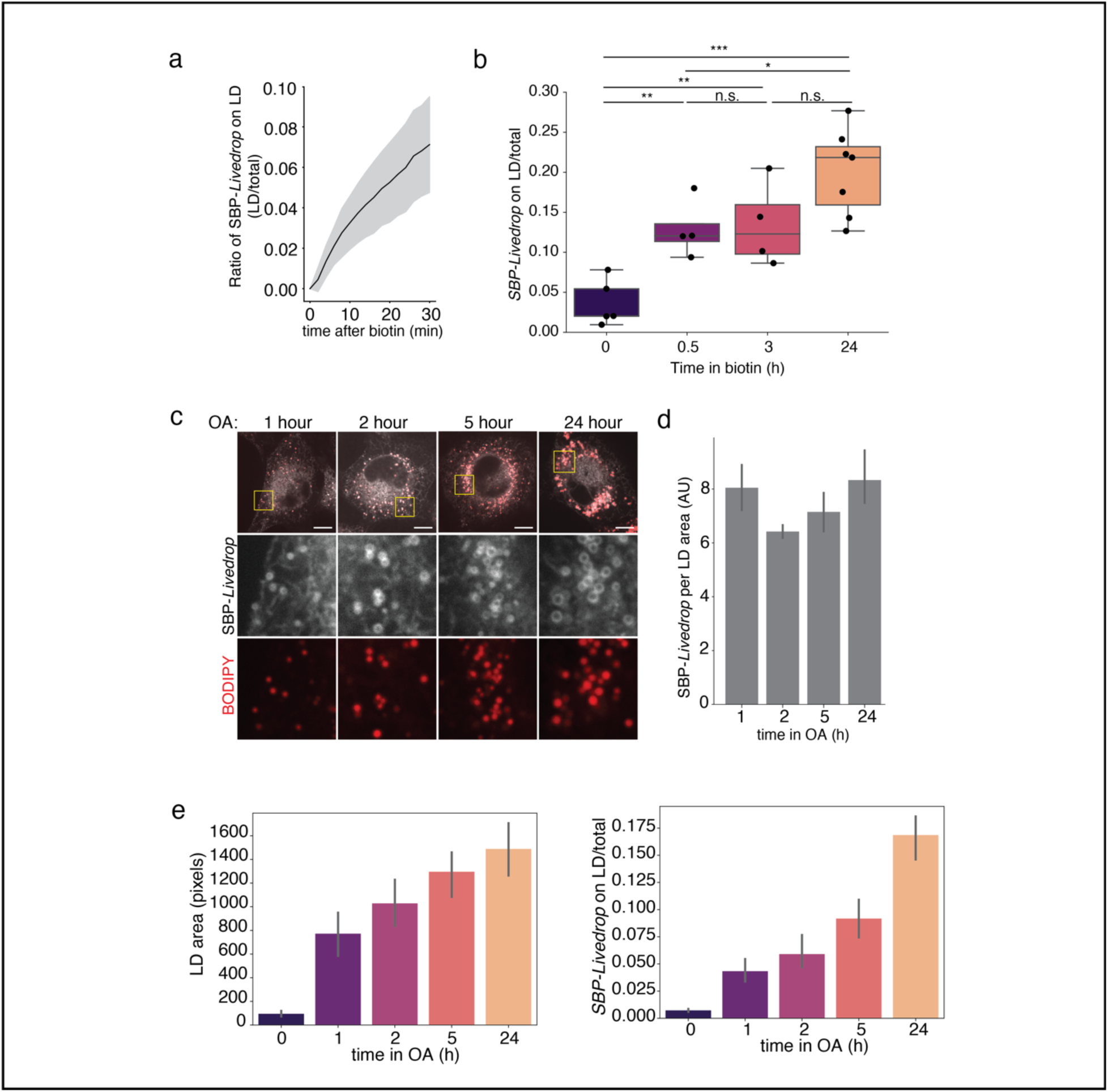
Synchronization of ER-to-LD protein transport using the RUSH system. **a.** Quantification of SBP-LiveDrop targeting kinetics after biotin-induced release. Targeting ratio was calculated as the intensity of SBP-LiveDrop within the BODIPY-stained LD mask normalized to total cellular SBP-LiveDrop intensity and baseline (pre-release) levels. *N* = 35. **b.** SBP-LiveDrop targeting ratio to LDs measured at multiple time points post-biotin release. Statistical significance was assessed by unpaired t-tests with p-values indicated for comparisons between time points. *P* values: 0.004 (0 vs 0.5 h), 0.009 (0 vs 3 h), 0.0001 (0 vs 24 h), 0.86 (0.5 vs 3 h), 0.04 (0.5 vs 24 h), 0.08 (3 vs 24 h). **c.** Representative confocal images showing SBP-LiveDrop localization to nascent and mature LDs after biotin release. Cells were pretreated with 250 µM oleic acid for varying durations to induce LD formation, followed by 30-min biotin release prior to imaging. LDs were stained with 1 µM BODIPY 493/503. Scale bar: 10 µm. **d.** Quantification of SBP-LiveDrop intensity normalized to LD area, calculated by dividing the targeting ratio by the total BODIPY mask area per image, reflecting SBP-LiveDrop density on LD surfaces after 30 min biotin release. **e.** Total LD area (left) and SBP-LiveDrop targeting ratio (right) quantified after 30 min biotin release in cells incubated with 250 µM oleic acid for increasing durations.

**Extended Data Fig. 6.**
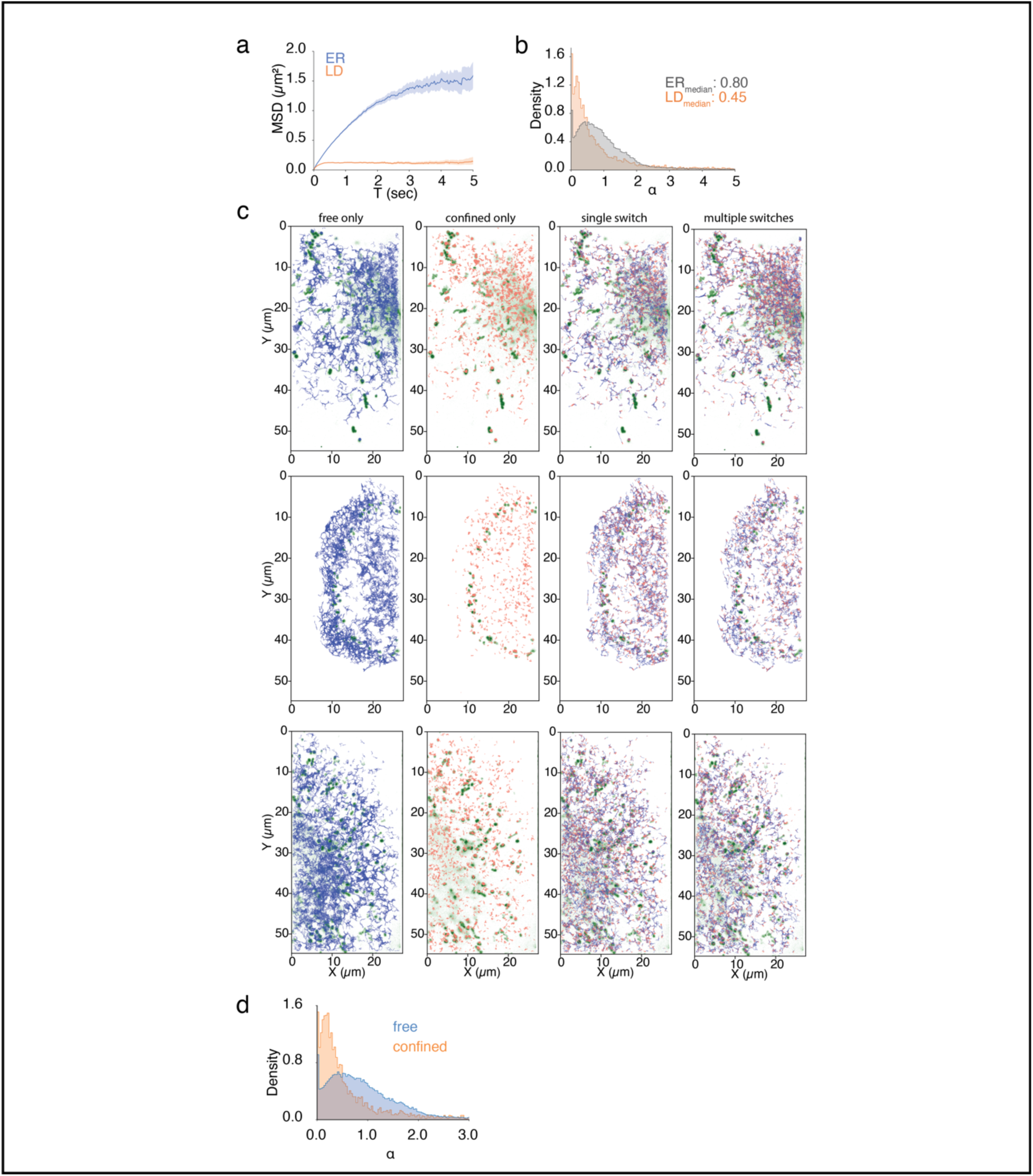
Classification of single-molecule tracks, based on motion states. **a.** Mean-square displacement (MSD) curves of tracks localized to the ER and LD, recorded by HILO imaging. **b.** Distribution of anomalous diffusion exponent (α) for ER-localized (blue) and LD-localized (orange) tracks. Median α values are indicated. **c.** Representative images showing motion classes of SBP-LiveDrop tracks. Tracks were classified using the DeepSPT diffusion pipeline, which analyzes changes in α within individual tracks. Four motion classes are defined: free motion, confined motion, single switch (one transition between free and confined), and multiple switches (multiple transitions). **d.** Distribution of α values for tracks classified as free (blue) and confined (orange).

**Extended Data Fig. 7.**
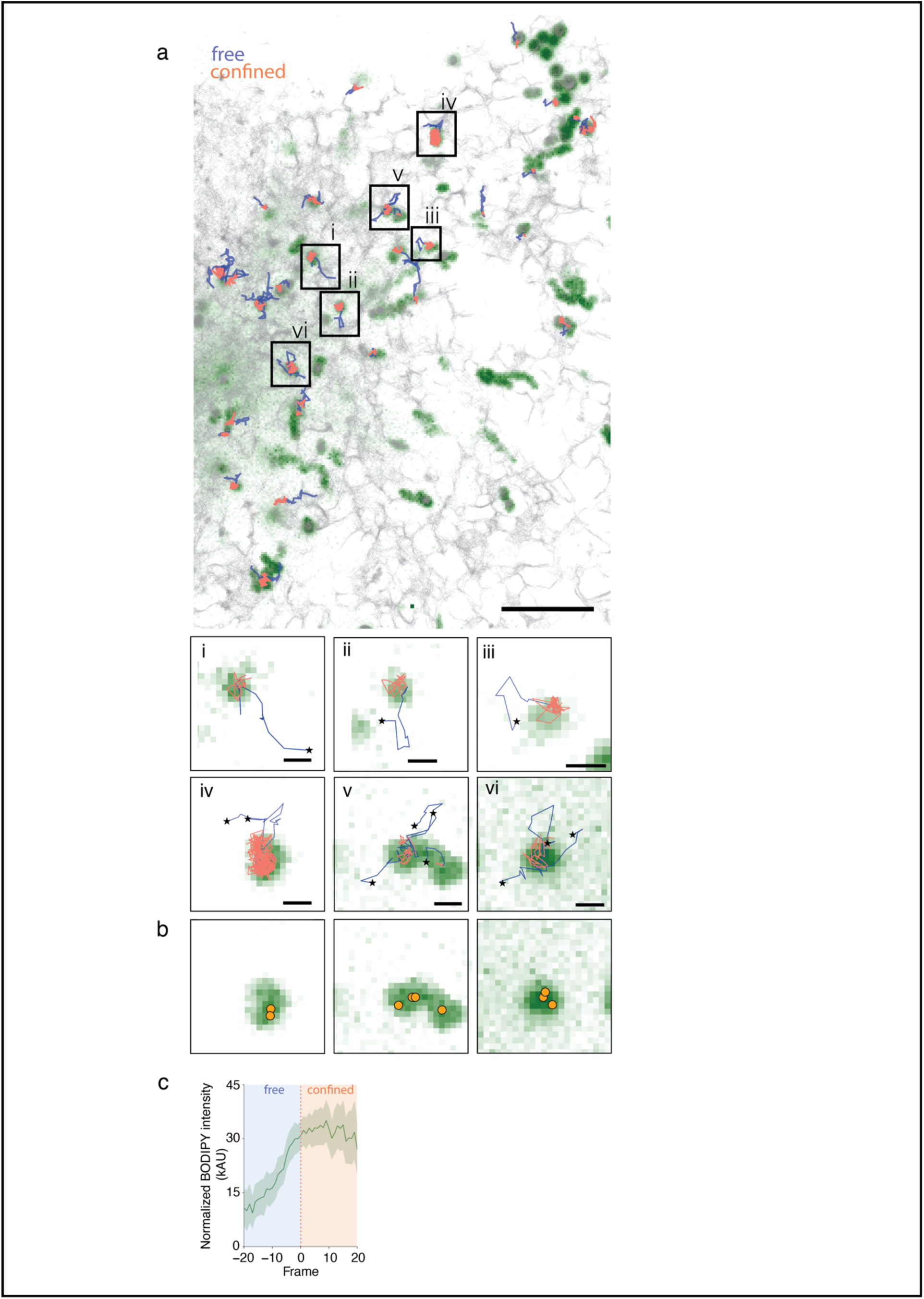
Protein entry onto the LD surface through ER-LD contact sites. **a.** Single-molecule tracks of SBP-*LiveDrop* 8 minutes after biotin release. Cells were treated with 250 µM oleic acid for 3 hours prior to release and labeled with 80 pM JFX554 for HILO imaging. Tracks transition from free motion (blue) to confined motion (orange) as proteins move from the ER membrane to the LD surface. All recorded tracks are shown in light grey. LDs are labeled with BODIPY 493/503 (green). Scale bar: 10 µm. Insets (bottom) highlight representative ER-to-LD trajectories, colored by motion state (blue: free, orange: confined); track start points are marked with stars. Scale bar: 500 nm. **b.** Overlay of coordinates where tracks switch from free to confined motion (from panel a) plotted over the LD channel (green). **c.** Mean BODIPY intensity along ER-to-LD tracks, aligned to the frame of motion-state switch (frame 0), with standard deviation shown. The increase in BODIPY signal after frame 0 indicates protein entry onto the LD surface. *N* = 219.

**Extended Data Fig. 8.**
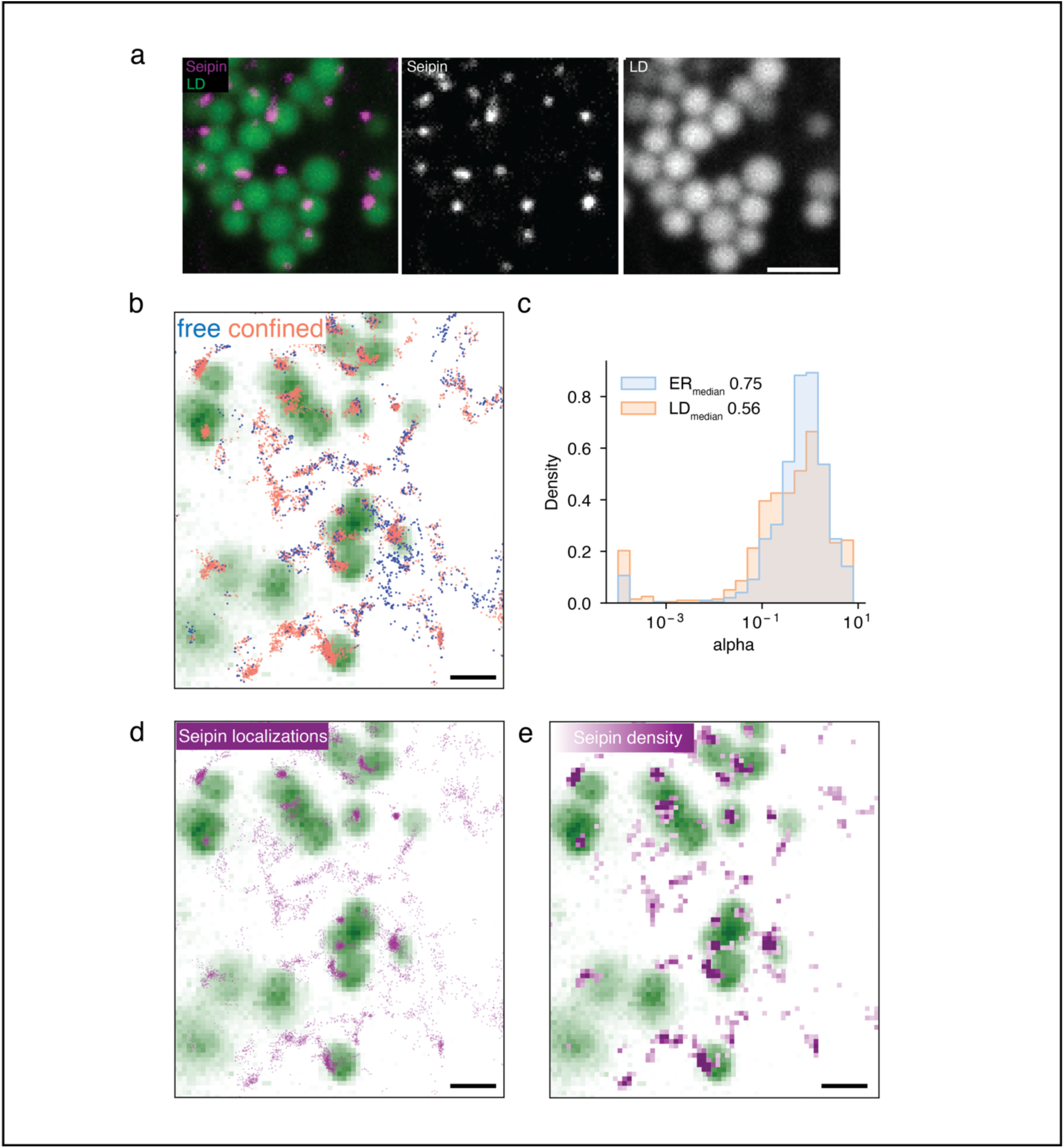
Spatial dynamics of endogenous Seipin at ER-LD contact sites. **a.** Representative spinning disk confocal images of endogenous Seipin-sfGFP in cells incubated overnight with 250 µM oleic acid and stained with LipidTOX Deep Red to label LDs. Scale bar: 2 µm. **b.** Single-molecule localizations of endogenously tagged Seipin, colored by motion class. Cells were transiently transfected with Halo-tagged Plin3 to label LDs, incubated overnight with 250 µM oleic acid, and Plin3 was labeled with JF554X prior to imaging. **c.** Distribution of the anomalous diffusion exponent (α) for Seipin tracks localized to the ER (blue) and LDs (orange). Median α values are indicated. **d.** Spatial plotting of Seipin single-molecule positions from individual complexes over the LD channel, showing enrichment of Seipin near LDs. Scale bar: 2 µm. **e.** Positional density map of Seipin localizations marking ER-LD contact sites. Density is calculated as the frequency of Seipin detections per pixel and represented as a colormap; darker magenta indicates higher density (>25 positions). Scale bar: 2 µm.

**Extended Data Fig. 9.**
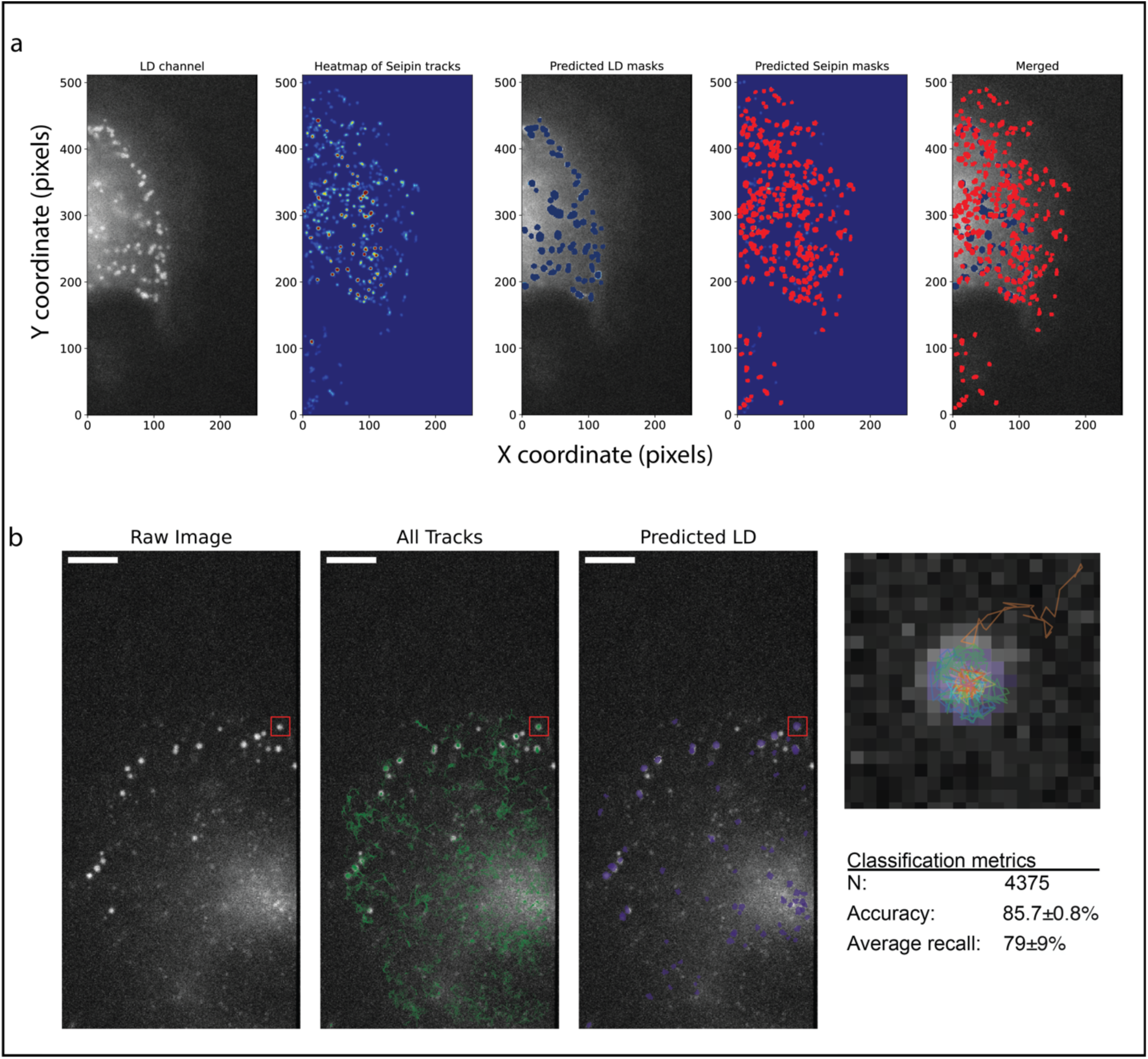
LD mask prediction using motion characteristics of *SBP-LiveDrop* tracks. **a.** From left to right: HILO images of LD labelled with BODIPY. Heatmap of seipin localization density. HILO images of LD labelled with BODIPY overlaid with LD segmentation. Heatmap of seipin with identified seipin hotspots overlaid. HILO images of LD labelled with BODIPY overlaid with LD mask segmentations and seipin hotspots. **b.** HILO images of LD labelled with BODIPY (left). SBP-*LiveDrop* tracks overlaid (middle). Predicted LD mask based on motion characteristics of SBP-*LiveDrop* (right, see Methods). Zoom-in: LD with SBP-*LiveDrop* tracks classified as LD associating overlaid exemplifying the identification of LD using SBP-*LiveDrop* tracks. Table: Classification metrics for the prediction of LD-localized trajectories to be used for identification of LD localization from SBP-*LiveDrop* tracks alone. Metrics are evaluated by Leave-one-out cross-validation by withholding a single SPT movie as test set and utilizing the remaining for training. N signifies the total number of tracks evaluated in the cross-validation scheme. Accuracy and recall quantify the classification performance across the cross-validation.

**Extended Data Fig. 10.**
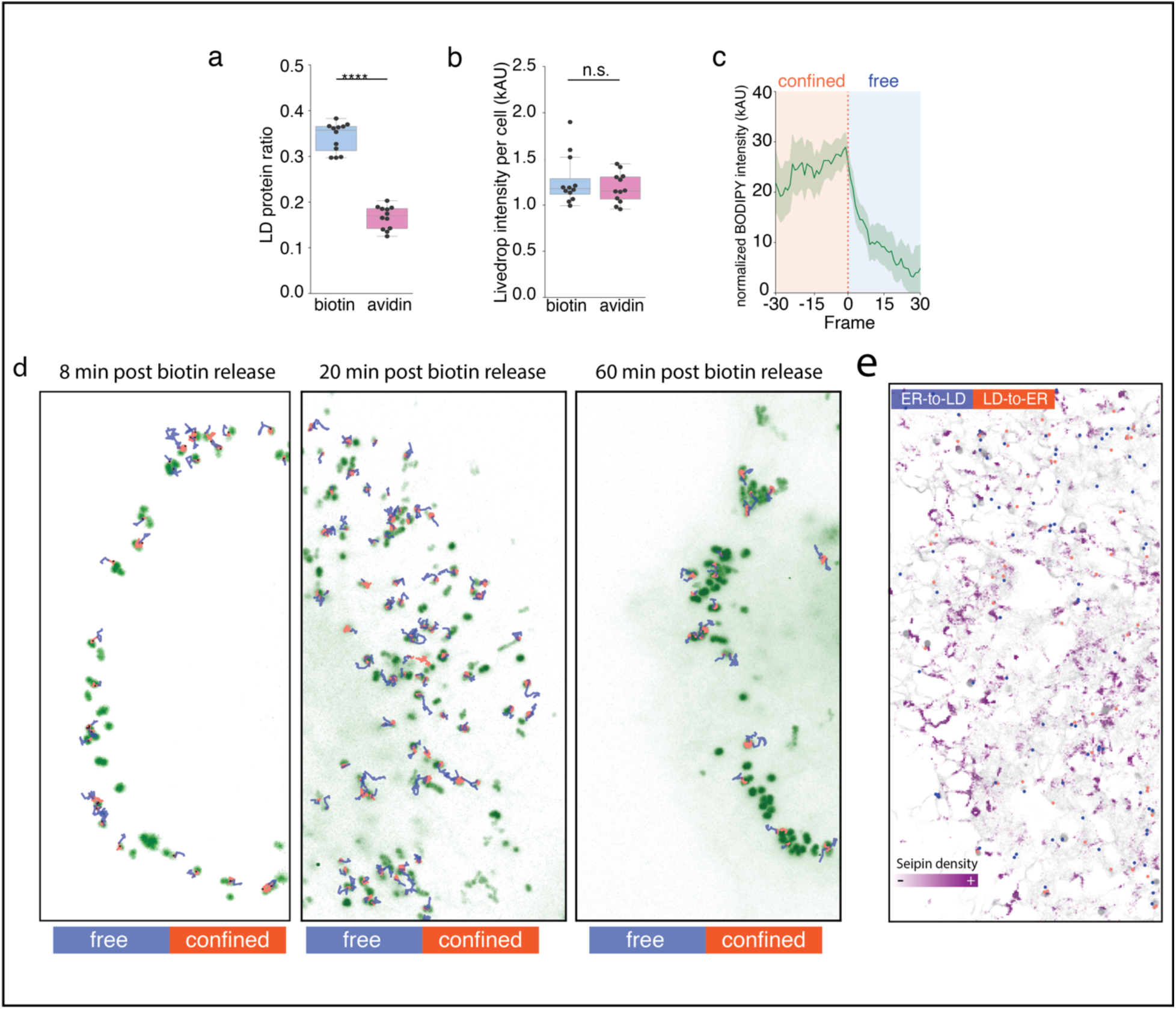
Bidirectional trafficking of *SBP-LiveDrop* single molecules. **a.** Quantification of *SBP-LiveDrop* targeting ratio. An unpaired t-test was performed to compare biotin and avidin samples. *P* value: 5.3×10^−10^. **b.** Quantification of total *SBP-LiveDrop* signal in cells. An unpaired t-test was performed to compare biotin and avidin samples. P value: 0.86. **c.** BODIPY signal intensity along single-molecule tracks that exited LDs. The switch from confined-to-free motion is indicated as Frame 0. Mean values are shown with standard deviation. *N* = 331. **d.** Examples of SBP-*LiveDrop* single-molecule tracks entering and exiting LDs at different stages of biotin release. Tracks are colored, based on motion properties. Blue: free; orange: confined. **e.** Motion switch coordinates of ER-to-LD (blue) and LD-to-ER (orange) single-molecule tracks overlaid onto a Seipin density map. All single-molecule tracks were colored in light gray.

